# Effects of age on the response to spinal cord injury: optimizing the larval zebrafish model

**DOI:** 10.1101/2023.05.18.541337

**Authors:** Whitney J. Walker, Kirsten L. Underwood, Patrick I. Garrett, Kathryn B. Lorbacher, Shannon M. Linch, Thomas P. Rynes, Chloe Sloop, Karen Mruk

**Author notes:** Address correspondence to:Dr. Karen Mruk, East Carolina University Brody School of Medicine, Department of Pharmacology and Toxicology 600 Moye Boulevard, Greenville, NC 27834. These authors contributed equally to this work (Kirsten L. Underwood, Patrick I. Garrett).

## Abstract

Zebrafish are an increasingly popular model to study regeneration after spinal cord injury (SCI). The transparency of larval zebrafish makes them ideal to study cellular processes in real time. Standardized approaches, including age at the time of injury, are not readily available making comparisons of the results with other models challenging. In this study, we systematically examined the response to spinal cord transection of larval zebrafish at three different larval ages (3-, 5-, or 7-days post fertilization (dpf)) to determine whether the developmental complexity of the larvae affects the overall response to SCI. We then used imaging and behavioral analysis to evaluate whether differences existed based on the age of injury. Injury led to increased expression of cytokines associated with the immune response; however, we found that the timing of specific inflammatory markers changed with the age of the injury. We also observed changes in glial and axonal bridging with age. Young larvae (3 dpf) were better able to regenerate axons independent of the glial bridge, unlike older larvae (7 dpf), consistent with results seen in adult zebrafish. Finally, locomotor experiments demonstrated that some swimming behavior occurs independent of glial bridge formation, further highlighting the need for standardization of this model and functional recovery assays. Overall, we found differences based on the age of transection in larval zebrafish, underlining the importance of considering age when designing experiments aimed at understanding regeneration.

## INTRODUCTION

Spinal cord injury (SCI) is a devastating event that leads to permanent loss of function in mammals. The zebrafish central nervous system (CNS) shares many organizational, cellular and molecular pathways with mammals including myelinated axons (Brösamle and Halpern, 2002) and use of the same neurotransmitters (Panula et al., 2006). Despite these similarities, regeneration and locomotor recovery occur in zebrafish even after complete transection of the spinal cord (van Raamsdonk et al., 1998). Understanding the similarities and differences between regenerative zebrafish and non-regenerative mammals could be pivotal for identifying new therapeutic targets and strategies to induce functional recovery in humans.

Adult zebrafish retain their regenerative ability throughout their lifetime, making them an obvious choice to compare with mammalian models. However, adult zebrafish take weeks– months to recover swim function (Vajn et al., 2014) and require complicated surgical procedures (Fang et al., 2012), which preclude the use of this model from many laboratories with smaller facilities that cannot support the animal number or specialized techniques required for adult zebrafish studies. Furthermore, adult zebrafish are not naturally transparent, limiting the use of real-time imaging of cellular processes during regeneration.

Larval zebrafish offer many experimental advantages over adult zebrafish to study SCI (Alper and Dorsky, 2022). Notably, larval zebrafish offer a transparent model to study regeneration in real time. In addition, the larvae’s small size permits individual animals to be tracked from injury to regeneration. Many processes are conserved between larval and adult zebrafish. Studies demonstrated that innate immune cells infiltrate the injury site in both larvae (Cavone et al., 2021; Gollmann-Tepeköylü et al., 2020; Tsarouchas et al., 2018) and adults (Hui et al., 2010; Hui et al., 2014) and are required for regeneration. SCI in both larvae (Ohnmacht et al., 2016) and adults (Reimer et al., 2013) leads to regeneration of motor neurons, which is enhanced through dopamine agonism. Lastly, both larval and adult zebrafish form a glial bridge across the injury site (Briona and Dorsky, 2014a; Goldshmit et al., 2012; Klatt Shaw et al., 2021; Matsuoka et al., 2016; Mokalled et al., 2016) which may promote axon growth after SCI. Although each of these studies highlights the utility of the larval model, differences in the age used for experiments limits the interpretation of these studies when compared with adults or mammals.

Embryonic development is classically defined as ending at the time of hatching, 3-4 days post fertilization (dpf) (Kimmel et al., 1995; Parichy et al., 2009) with larval development continuing for approximately six weeks. Often laboratories use late embryonic and early larval stages due to ease and ethical considerations (Tsarouchas et al., 2018) with some studies looking at larvae as old as 10 dpf (Hossainian et al., 2022). Furthermore, neurogenesis of primary motor neurons is greatly reduced at 2 dpf (Reimer et al., 2013) and nascent myelin is observed 3-4 dpf (Bin and Lyons, 2016; Kirby et al., 2006; Park et al., 2002; Preston and Macklin, 2015). Although injury-dependent mechanisms have been identified in these earlier ages, these studies can be limiting as additional factors such as mature synaptic connections and myelin debris are largely absent in zebrafish injured this young. We sought to determine whether larvae transected at different ages demonstrated differences in the recovery process. We transected the spinal cord of larval zebrafish at 3 dpf, 5 dpf, and 7 dpf and characterized the subsequent cellular events and locomotor recovery. We show that some secondary injury mechanisms and subsequent regeneration changes with age. Taken together, our data suggests that age should be considered when designing larval zebrafish SCI experiments.

## RESULTS

### Age-dependent injury response in zebrafish larvae

Many larval zebrafish SCI experiments use a mechanical transection above the cloaca to create the SCI. As other organ systems are still maturing at larval zebrafish ages (Brown et al., 2016), we first sought to determine whether the damage to the skin and muscle from a mechanical transection may be causing changes to the developing zebrafish. Therefore, using bright field microscopy, we first determined whether mechanical transection of the spinal cord affected basic larval biology after SCI. We transected *Tg(gfap:EGFP)* zebrafish larvae at 3 dpf, 5 dpf, or 7 dpf and used the signal from green fluorescent protein (EGFP) to confirm the presence of a complete glial transection. We chose these ages as they correlate with the end of embryonic stage (Parichy et al., 2009), development of feeding (Hernandez et al., 2018), and mineralization of the centra (Bensimon-Brito et al., 2012). Both age-matched control larvae that were not injured and fully transected larvae were recorded daily. We first looked at normal physiological development. In zebrafish, the inner most lining of the swim bladder is formed by 3 dpf, with inflation of the chamber occurring at approximately 4.5 dpf (Winata et al., 2009). After transection, we found that larvae injured at 3 dpf failed to properly inflate swim bladders as late as five days after injury (8 dpf, 22% compared with 96% for control larvae) (Fig. S1A). To ensure this was due to the SCI and not an artifact of the *Tg(gfap:EGFP)* line, we also monitored the inflation of the swim bladder in *Tg(olig2:dsRed*) larvae and similarly found that swim bladder inflation was reduced in transected larvae (8 dpf, 12% vs. 100% for control larvae) (Fig S1B) indicating that injury affected normal swim bladder development. We routinely observed normal swim bladders after transection of 5 dpf and 7 dpf larvae. We also measured the heart rate of transected larvae and tracked individual fish over time to determine whether survival of larvae could be predicted (Fig. S1C). Larvae injured at 3 dpf, showed a slight, but not statistically significant increase in heart rate at 1 day post injury (dpi) which stayed elevated through 3 dpi. In contrast, larvae injured at 5 dpf showed a decrease in heart rate on the day after injury with larvae injured at 7 dpf showing no differences in heart rate across all ages (Fig. S1C). For all ages, individual larvae with greatly reduced blood flow were most likely to die over the course of the week. Given that the small differences observed in heart rate across ages were not statistically significant, additional cellular mechanisms may be leading to previously reported mortality rates observed after a mechanical injury in zebrafish larvae (Bhatt et al., 2004).

We next used mechanical transection to focus on the larval zebrafish response to SCI. There are two mechanisms of damage after SCI: a primary mechanical injury and a secondary injury mediated by many processes including continued apoptosis and inflammation. In mammals, apoptosis can occur hours after injury and persist for weeks (Mizuno et al., 1998). To determine the amount of apoptotic cells in the spinal cord after injury, we incubated live *Tg(olig2:dsRed)* larvae with acridine orange from 12 hours post injury (hpi) – 5 dpi. Using the dsRed signal to exclude acridine orange signal from outside of the CNS, we observed day-to-day differences between the ages of injury. For larvae injured at 3 dpf or 5 dpf, we found a significant increase in the total number of apoptotic cells at 12 hpi and 24 hpi (Fig. 1A-D) compared to controls. In these larvae, apoptosis began to significantly resolve at 72 hpi (p value for 12 hpi vs 72 hpi = 0.001 and 0.0092 respectively). In contrast, there was no reduction in the number of apoptotic cells among larvae transected at 7 dpf (p value for 12 hpi vs 120 hpi >0.9999) (Fig. 1E-F). Resolution of apoptosis followed an exponential decay for 3 dpf and 5 dpf with older larvae (7 dpf) have a sustained injury process (Fig. 1G).

**Fig. 1.**
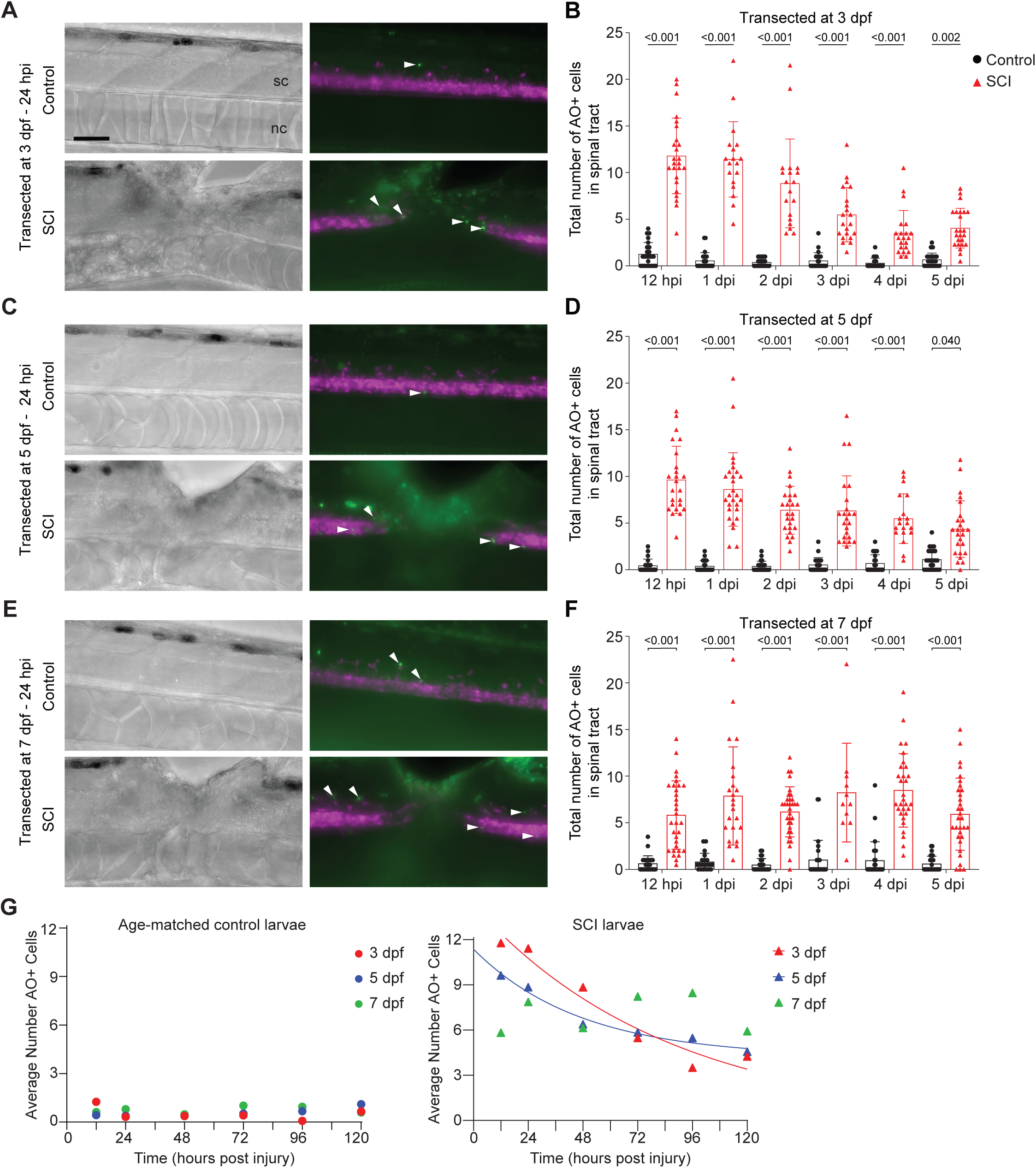
Resolution of cell death varies with age of transection. Representative micrographs from *Tg(olig2:dsRed)* larvae incubated in acridine orange (AO) 24 hours post injury (hpi) are shown for (**A**) 3 dpf, (**C**) 5 dpf, and (**E**) 7 dpf larvae, with green signal indicating cell death. Positive cells (white arrowheads) were counted in the region of the spinal cord, determined by dsRed signal. Larvae orientation: lateral view, anterior left. Scale bar = 50 µm. Quantification of apoptotic cells in larvae transected at (**B**) 3 dpf, (**D**) 5 dpf, or (**F**) 7 dpf. The total number of cells in individual fish are plotted as determined by two independent researchers. Mean ± SD is shown. (**G**) Average number of apoptotic cells plotted over time for age-matched uninjured and transected larvae. Lines indicate a least squares fit of a one phase exponential decay. Statistics for the observed phenotypes are in Table S1.

Given the sustained levels of apoptosis in older larvae, we next asked whether the inflammatory response was also extended in older fish. We used quantitative RT-PCR (qRT-PCR) to analyze the levels of cytokine expression after injury in these three ages compared to their non-transected controls (Fig. 2). In pooled larvae transected at 3 dpf, expression of the pro-inflammatory cytokine *il-1*β was high initially after injury (∼7-fold on day of injury) but did not peak in larvae injured at 7 dpf until 4 dpi (∼10-fold change) (Fig. 2A). Similarly, *tgf-*β*1* and *tgf-* β*3* were upregulated the day of injury in larvae transected at 3 dpf or 5 dpf, respectively, consistent with previous 3 dpf transection studies (Fig. 2B-C). However, we measured only a 2-3 fold increase in *tgf-*β*1* levels and no change in *tgf-*β*3* expression in larvae transected at 7 dpf. In contrast, the pro-inflammatory cytokine, *tnf-*α, was upregulated 2 days after injury in 7 dpf larvae, with little expression in the first days after injury in larvae transected at 3 dpf or 5 dpf (Fig. 2D).

**Figure 2.**
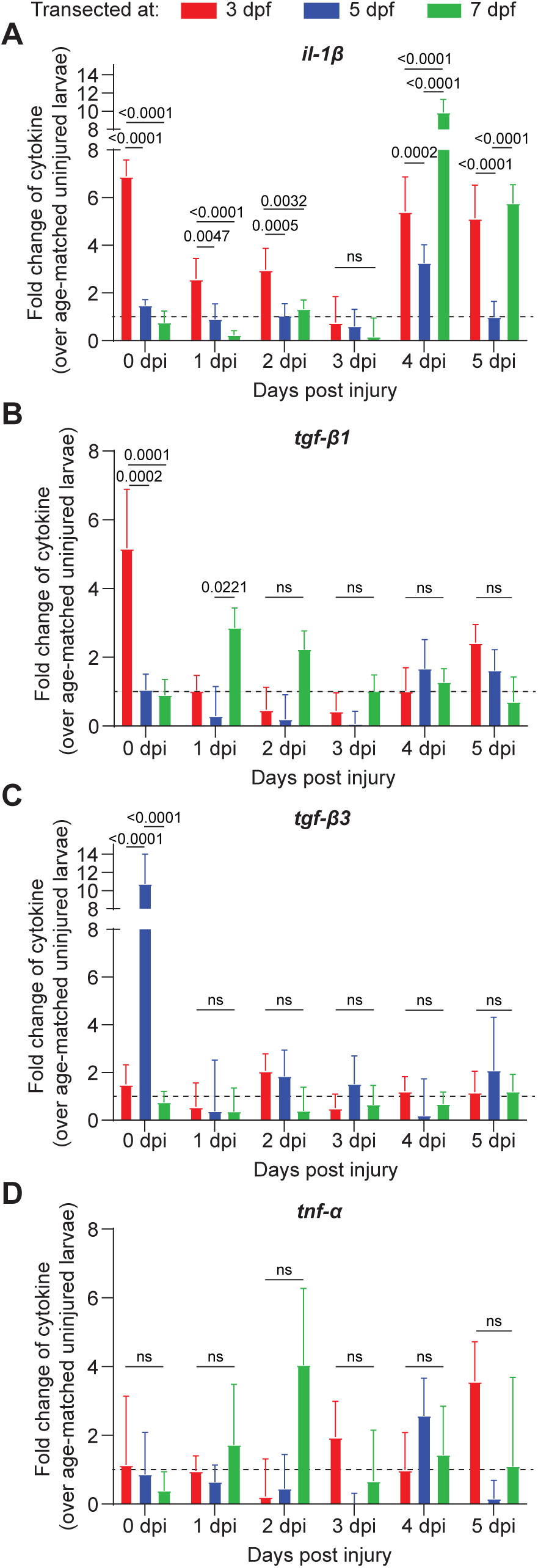
Cytokine expression changes with the age of transection in zebrafish larvae. *Tg(gfap:EGFP)* larvae were transected at 3 dpf, 5 dpf or 7 dpf and collected each day post injury (dpi). The expression fold change of (**A**) *il-1*β (**B**) *tgf-*β*1*, (**C**) *tgf-*β*3*, and (**D**) tnf-α over age-matched uninjured larvae was calculated. Larvae transected at 3 dpf transection in red, 5 dpf in blue and 7 dpf in green. Mean ± SEM of at least three technical replicates from pooled samples containing five different larvae are plotted. Two-way ANOVA with Tukey multiple comparisons test for each day were compared. Adjusted p value is indicated. ns: p>0.100

### Glial and axons cross the site of injury at different rates with age

A hallmark of the adult zebrafish response to spinal cord injury is the formation of a glial bridge (Goldshmit et al., 2012; Mokalled et al., 2016). Therefore, we next determined whether larval zebrafish faithfully recapitulated glial recovery seen in adult zebrafish. We generated heterozygous *Tg(elavl3:mCherry-CAAX)* zebrafish and crossed them with *Tg(gfap:EGFP)* zebrafish to generate larvae which contained labeled axons and radial glia. Both the sensory and motor tracts are clearly labeled in these larvae. The advantage of this line permits sequential imaging of the same transected larvae over the course of the experiment instead of relying on pooled fixed larvae for immunohistochemistry. We analyzed the formation of a radial glial bridge and ability of axons to cross the site of injury in larvae that had a complete spinal cord transection at 3 dpf, 5 dpf, or 7 dpf using confocal microscopy (Fig. 3). The radial glia in all transected larvae exhibited the well-characterized morphological change (Fig. 3A-C, *left panel*), of becoming more elongated (Briona and Dorsky, 2014a; Kim et al., 2008; Matsuoka et al., 2016). We used GFP pixel intensity to objectively quantify the injury size (Fig. SF2-4). For all ages, the average transection width was between 100-200 microns (Fig. 3A-C). Analysis of individual fish demonstrated that larvae transected at 3 dpf had a more variable glial bridging pattern with some fish bridging within a day after injury (Fig. SF2). In contrast, in older larvae, the width of the glial injury began to narrow before bridging was observed (Fig. SF3, 4).

**Figure 3.**
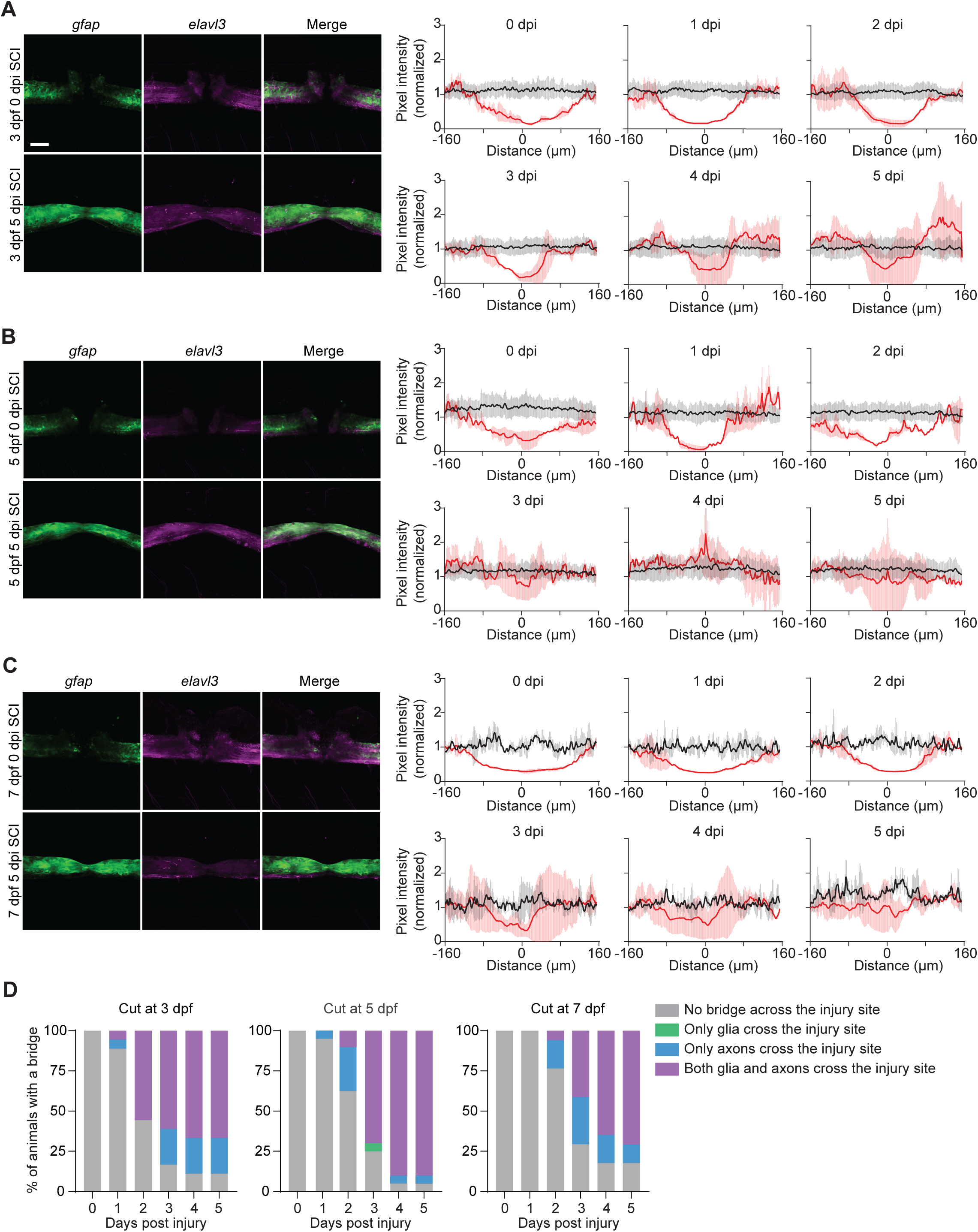
All ages of larvae form radial glia and axonal bridges across the injury site. *Tg(gfap:EGFP; elavl3:mCherry-CAAX)* larvae were transected and followed individually for 5 days post injury (5 dpi). *Left panels:* Representative micrographs from a maximal intensity projection of the spinal cord. Individual single larvae were tracked daily. The day of injury (0 dpi) and end of experiment (5 dpi) for larvae transected at: (**A**) 3 dpf, (**B**) 5 dpf, (**C**) 7 dpf are shown. Larvae orientation: lateral view, anterior left. Scale bar: 50 µm. *Right panels:* Quantification of mean pixel intensity (solid line) and standard deviation (shading) for control (black) and transected (red) larvae graphed approximately every 10 µm for clarity. Epicenter of the lesion denoted as 0 on the X-axis was determined by the lowest average normalized pixel intensity value on day of injury (0 dpi). Pixel intensity increases proximal to the lesion and persists through bridge formation indicating a local increase GFP signal. (**D**) Percent of animals with a bridge across the injury site was determined by complete tracks of fluorescence across the site of injury. 3 dpf: control: n = 24, SCI: n = 18; 5 dpf: control: n=26, SCI: n = 16; 7 dpf: control: n=21, SCI: n = 16 SCI, all tracked individually. Statistics for the observed phenotypes are in Table S2.

In adult zebrafish, cells accumulating at the edge of the lesion express low levels of Gfap. (Goldshmit et al., 2012). Therefore, to confirm that changes observed in the GFP intensity after injury was not due to an increase in *gfap* expression, we used qRT-PCR. We found that larval zebrafish transected at 3 dpf and 5 dpf had either reduced or unchanged expression levels of *gfap* until 4 dpi (Table 1). In contrast, older larvae showed relatively stable expression levels of *gfap,* suggesting that changes in pixel intensity observed is not just due to increased levels of GFP. One gene required for glial bridging in adult zebrafish is connective tissue growth factor a (*ctgfa*) (Mokalled et al., 2016). We also found that independent of transection age, all larvae upregulated *ctgfa* between 2-3 dpi which remained elevated even after glial bridging (Table 1). Taken together, the cellular accumulation of Gfap+ cells and timing of Gfap+ cells bridging the injury gap differs with age of injury.

**Table 1.** Relative fold changes of *gfap* and *ctgfa* after injury.*^a^*.

| Gene | Days post injury (dpi) | 3 dpf SCI | 5 dpf SCI | 7 dpf SCI |
| --- | --- | --- | --- | --- |
| *gfap* | 1 dpi | 0.65 ± 0.27 | 0.20 ± 0.70 | 1.8 ± 0.55 |
| *gfap* | 2 dpi | 0.47 ± 0.93 | 0.57 ± 0.33 | 0.89 ± 0.70 |
| *gfap* | 3 dpi | 0.75 ± 0.97 | 0.96 ± 1.02 | 1.5 ± 0.21 |
| *gfap* | 4 dpi | 3.0 ± 0.80 | 1.4 ± 0.72 | 1.0 ± 0.42 |
| *gfap* | 5 dpi | 0.47 ± 0.78 | 1.0 ± 0.56 | 1.2 ± 0.60 |
| *ctgfa* | 1 dpi | 0.15 ± 0.68 | 0.21 ± 2.05 | 0.65 ± 0.87 |
| *ctgfa* | 2 dpi | 0.34 ± 1.5 | 1.1 ± 0.67 | 0.86 ± 1.72 |
| *ctgfa* | 3 dpi | 0.59 ± 0.65 | 2.5 ± 0.61 | 2.8 ± 0.62 |
| *ctgfa* | 4 dpi | 3.48 ± 5.2 | 4.3 ± 0.38 | 4.0 ± 0.66 |
| *ctgfa* | 5 dpi | 0.67 ± 1.1 | 2.4 ± 0.55 | 6.9 ± 0.92 |

<sup>a</sup>Fold changes are presented as mean ± SEM.

We next examined whether glial bridging correlated with axonal growth across the injury site. Fifty percent of larvae transected at 3 dpf formed a glial bridge with axons crossing the site of injury within two days (Fig. 3D). As these larvae aged, we again observed that not all larvae formed a glial bridge (Fig. SF5); however, axonal growth continued such that 89% of larvae had axons that crossed the site of injury. Older larvae took longer for both glia and axons to cross the site of injury (3–4 days for 50% of larvae), with larvae cut at 7 dpf having the lowest percentage of axons crossing the site of injury (Fig. 3D).

### Recovery of free swim occurs independently of glial bridging, but large movements require both glial and axonal bridges

One aspect of complex behaviors in zebrafish is their ability to move at various speeds, which requires signals coming from the brain and local spinal cord circuitry (Berg et al., 2023; Kishore et al., 2014; Severi et al., 2014). Typically, researchers assay a single property such as total distance after SCI (de Sena-Tomas et al., 2024). Therefore, we first determined the recovery of total swim distance after a SCI in individually tracked 3 dpf, 5 dpf, and 7 dpf *Tg(GFAP:EGFP; elavl3:mCherry-CAAX)* larvae. After visual confirmation of a full transection by a blinded observer, we observed that larvae transected at 3 dpf recovered the most total swim behavior (Fig. 4A-C). In addition, 3 dpf age-match control zebrafish also increased their total swim distance over time, whereas 5 dpf and 7 dpf larvae did not. On the last day of the experiment, an independent observer imaged the zebrafish larvae and scored them for the presence of a bridge. For all ages, there was no difference in total distance in larvae independent of a glial and/or axonal bridge (Fig. 4A-C). We also measured the total number of movements and found an increase in the number of movements in 3 dpf larvae correlated with the increase in total distance (Fig. 4D-F).

**Figure 4.**
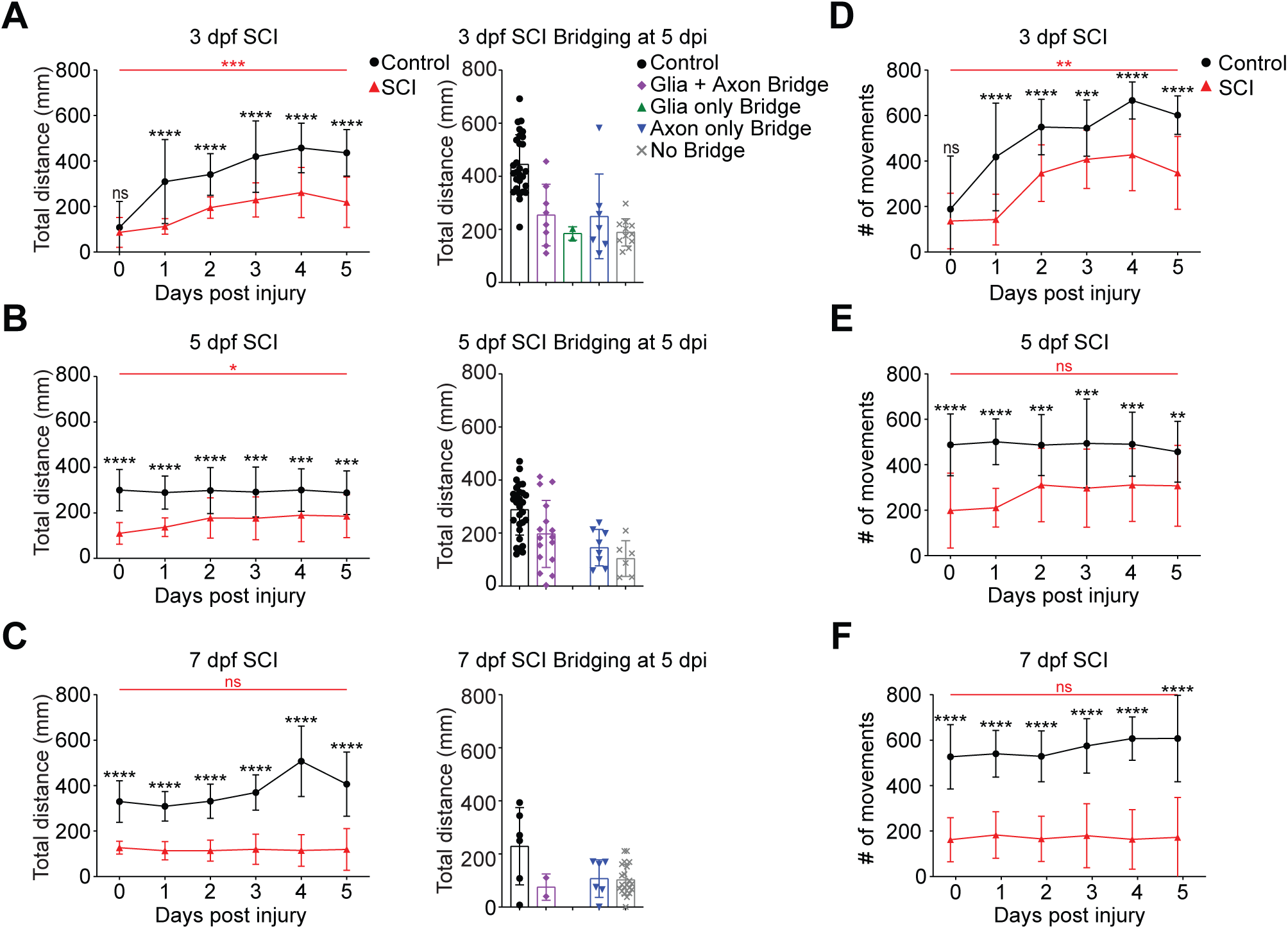
Recovery of total swim movement does not require glial and axonal bridging. *Tg(gfap:EGFP; elavl3:mCherry-CAAX)* larvae were transected and free swim followed individually. (**A-C**) *Left panels:* Total swim distance (mm) of control (black) vs transected larvae (red) generated from all swim speeds. Mean ± SD are shown. A two-way ANOVA using Tukey’s multiple comparison test was used to determine significance. *Right panels:* Quantification of total swim distance classified by bridging across the injury site at the end of the experiment (5 dpi). An experimenter blinded to the swim analysis imaged larvae for the presence of a bridge. Mean ± SD is shown with individual fish plotted. Statistics for the observed phenotypes are in Table S3. (**D-F**) Total number of swim movements of control (black) vs transected larvae (red) over the 10-min period of recording. Mean ± SD are shown. A two-way ANOVA using Tukey’s multiple comparison test was used to determine significance. For all experiments: 3 dpf: Control: n=28, SCI: n=28; 5 dpf: Control: n=29, SCI: n=30; 7 dpf: Control: 46, SCI: n=39 *p<0.05, **p<0.01, ***p<0.001, ****p<0.0001, ns: p≥0.2. Red text indicates comparisons between SCI 0 dpi and 5 dpi. Black text indicates comparisons between age- matched control and transected larvae.

Given that larvae can produce total distances through different swim durations and/or speed—which may not require a cellular bridge across the site of injury—we used video tracking to analyze swim distances occurring from twitch (0-2 mm/s), small (2-4 mm/s), and large (>4 mm/s) movements. Larvae injured at 3 dpf, 5 dpf, or 7 dpf were able to execute twitch movements regardless of injury with young larvae (3 dpf) generating more distance from twitch movements after a spinal transection than age-matched uninjured controls (Fig. 5A-C). For all ages, twitch movement was not dependent on glia and/or axons crossing the injury site (Fig. 5A- C). Looking at the number of twitch movements, we saw that 3 dpf fish increased the frequency of twitch movement over time independent of an injury, but the frequency of this movement was consistent across 5 dpf and 7 dpf larvae (Fig. 5D-F). As with twitch movement, 3 dpf larvae were able to generate distance from small movements (Fig. 7A, left) and the frequency of this movement increased as the larvae developed (Fig. 7D). In older larvae (5 dpf and 7 dpf), there was no appreciable change in distance from small movements or the frequency of small movements (Fig. 7B-C, E-F). Similar to twitch, all ages were capable of generating small movements without a glial bridge (Fig. 7A-C, right). In contrast, the distance from large movements increased over time for all three ages in transected larvae (Fig. 8A-C, left). While 3 dpf larvae were able to generate distance from large movements with just an axonal bridge, both a glial and axonal bridge was required in larvae transected at 5 dpf (Fig. 8A-C, right). Similarly, the amount of large movements increased during recovery in all three ages (Fig. 8D-F). Taken together, large swim movement best reports on recovery from SCI in zebrafish larvae. We repeated swim analysis on a second set of larvae that were monitored for a full week to determine whether behavior would recover with increased time (Fig. 9). Although more larvae had both a glial and axonal bridge across the injury site for all ages, we did not observe a substantial increase in swim behavior at 7 dpi compared to 5 dpi.

**Figure 5.**
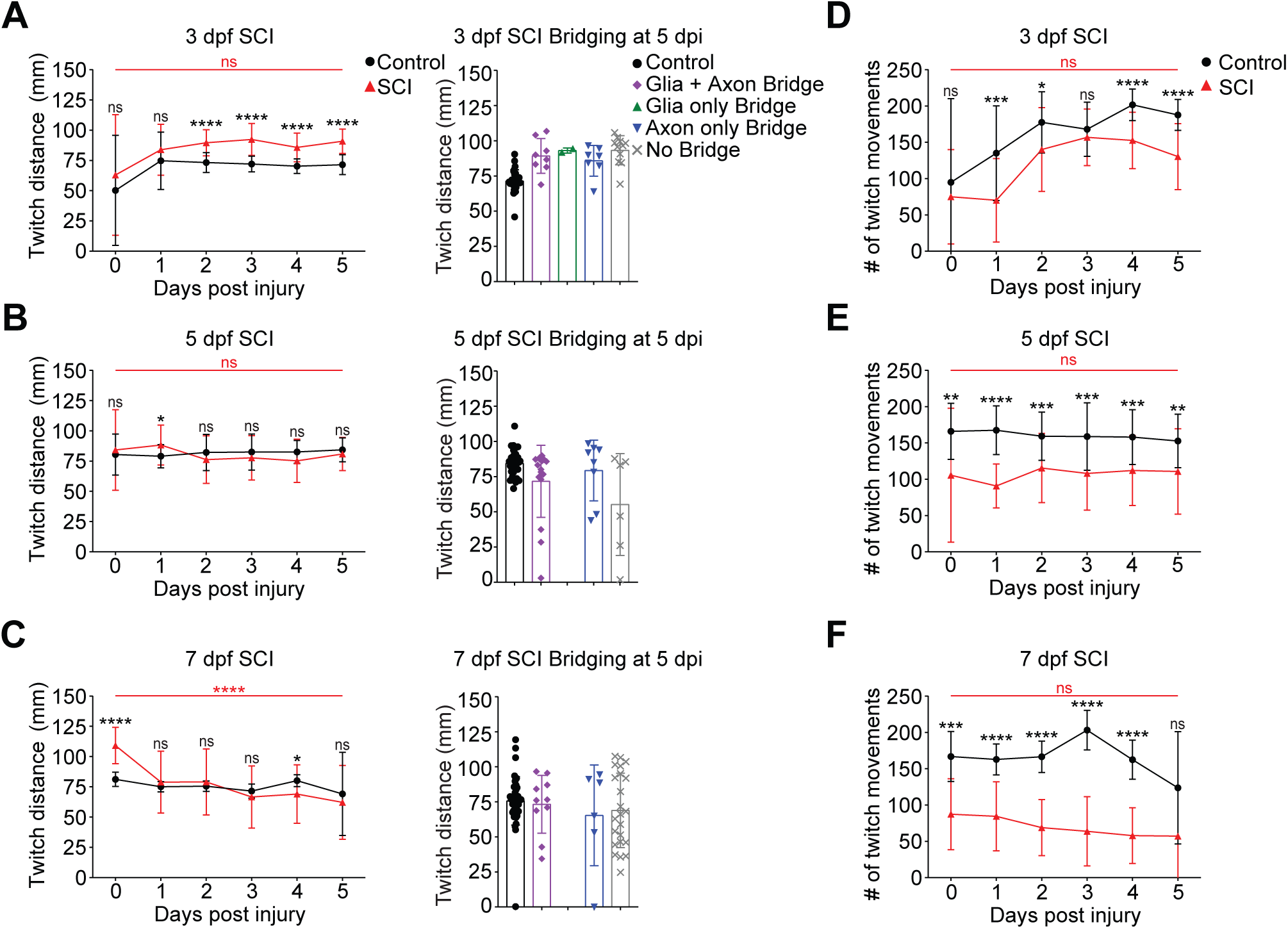
Injured larvae produce twitch movements after SCI without a glial or axonal bridge. (**A-C**) *Left panels:* Total swim distance of control (black) vs transected larvae (red) generated from twitch movements (swim speed: 0-2 mm/s). Mean ± SD are shown. A two-way ANOVA using Tukey’s multiple comparison test was used to determine significance. *Right panels:* Quantification of swim distance from twitch movements classified by bridging across the injury site at the end of the experiment (5 dpi). An experimenter blinded to the swim analysis imaged larvae for the presence of a bridge. Mean ± SD is shown with individual fish plotted. Statistics for the observed phenotypes are in Table S3. (**D-F**) Total number of twitch movements of control (black) vs transected larvae (red) over the 10-min period of recording. Mean ± SD are shown. A two-way ANOVA using Tukey’s multiple comparison test was used to determine significance. For all experiments: 3 dpf: Control: n=28, SCI: n=28; 5 dpf: Control: n=29, SCI: n=30; 7 dpf: Control: 46, SCI: n=39 *p<0.05, **p<0.01, ***p<0.001, ****p<0.0001, ns: p≥0.09. Red text indicates comparisons between SCI 0 dpi and 5 dpi. Black text indicates comparisons between age-matched control and transected larvae.

**Figure 6.**
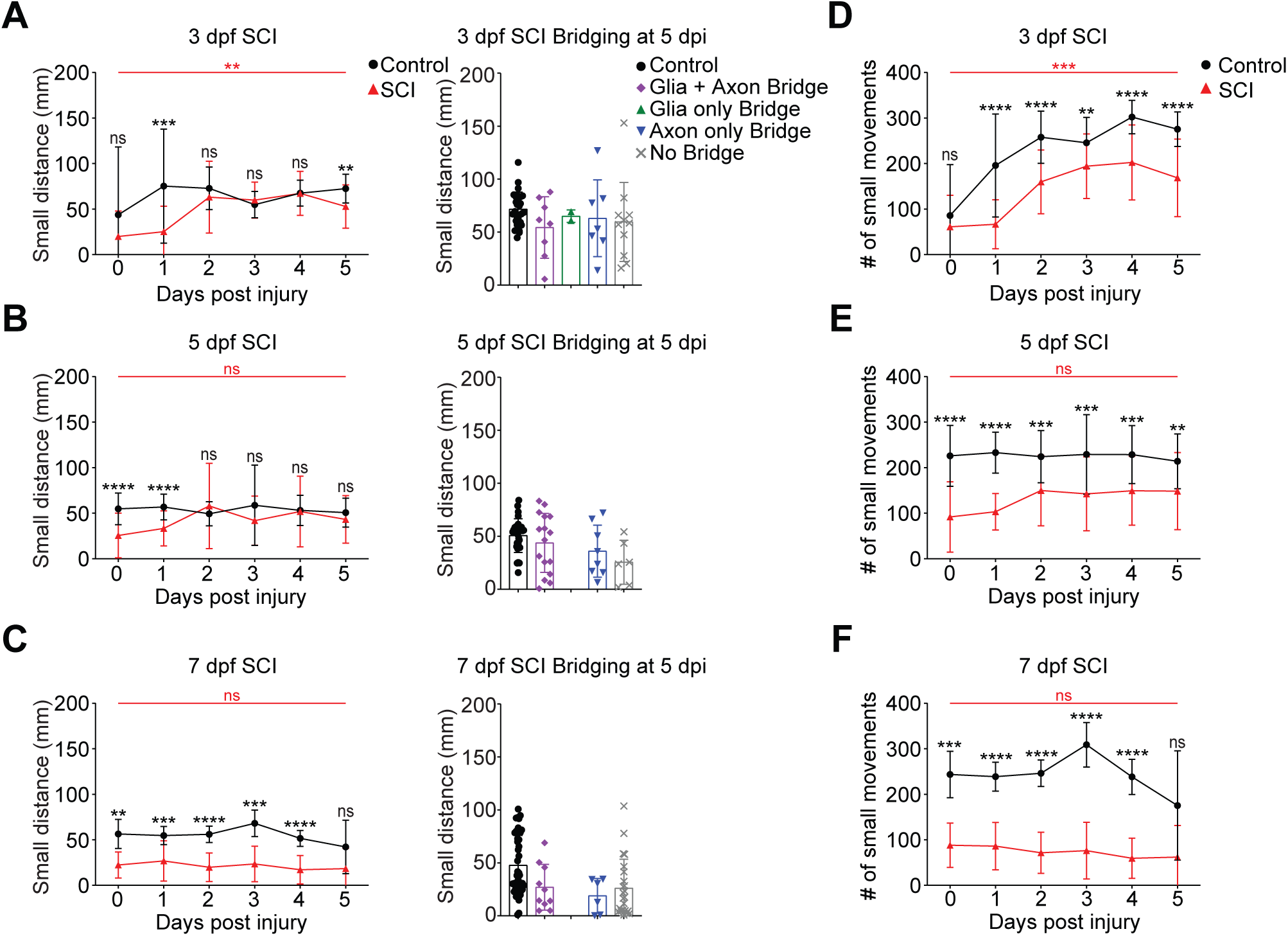
Injured larvae produce small movements after SCI without a glial or axonal bridge. (**A-C**) *Left panels:* Total swim distance of control (black) vs transected larvae (red) generated from small movements (swim speed: 2-4 mm/s). Mean ± SD are shown. A two-way ANOVA using Tukey’s multiple comparison test was used to determine significance. *Right panels:* Quantification of swim distance from small movements classified by bridging across the injury site at the end of the experiment (5 dpi). An experimenter blinded to the swim analysis imaged larvae for the presence of a bridge. Mean ± SD is shown with individual fish plotted. Statistics for the observed phenotypes are in Table S3. (**D-F**) Total number of small movements of control (black) vs transected larvae (red) over the 10-min period of recording. Mean ± SD are shown. A two-way ANOVA using Tukey’s multiple comparison test was used to determine significance. 3 dpf: Control: n=28, SCI: n=28; 5 dpf: Control: n=29, SCI: n=30; 7 dpf: Control: 46, SCI: n=39 *p<0.05, **p<0.01, ***p<0.001, ****p<0.0001, ns: p≥0.2. Red text indicates comparisons between SCI 0 dpi and 5 dpi. Black text indicates comparisons between age- matched control and transected larvae.

**Figure 7.**
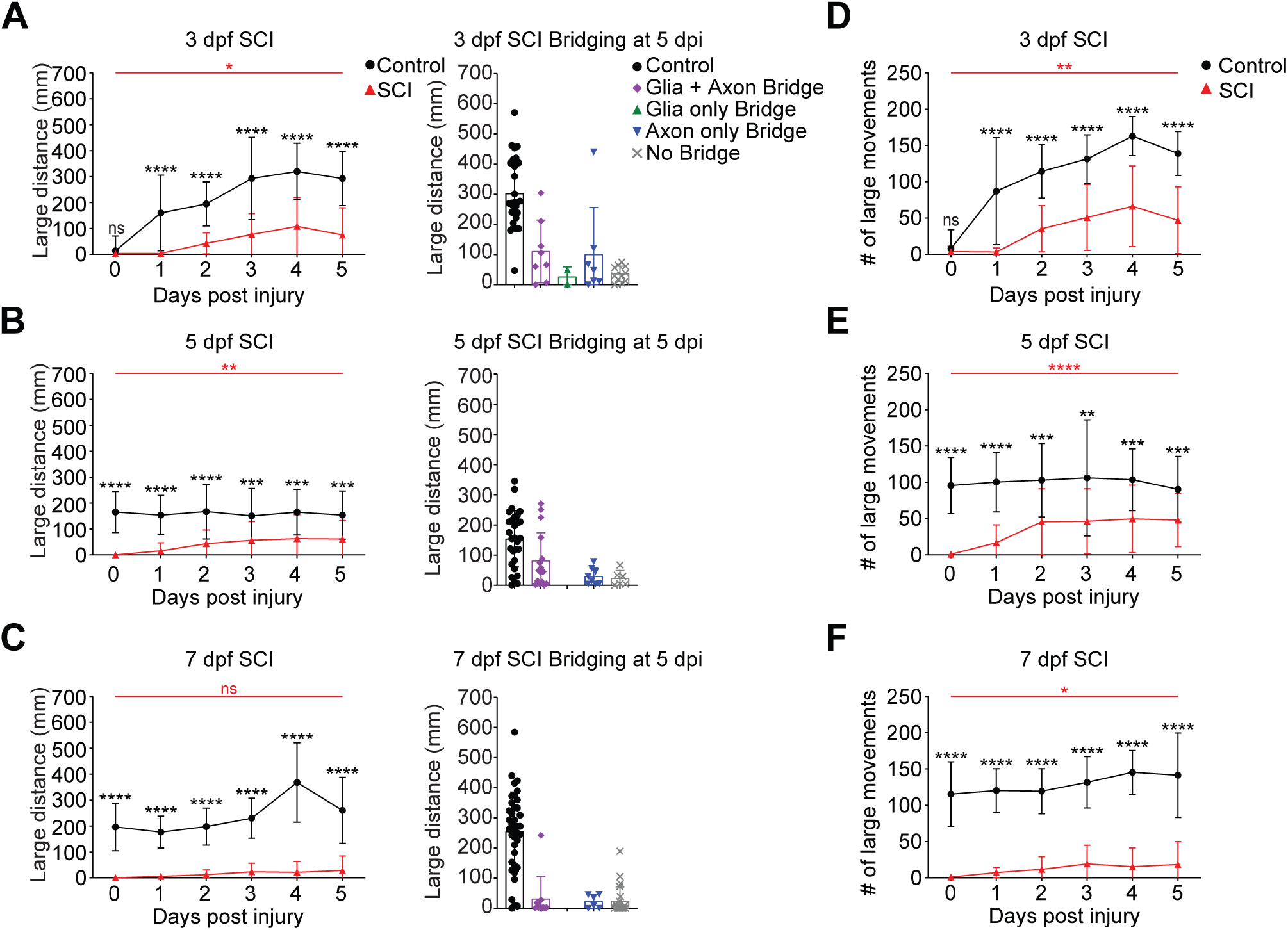
Injured larvae require glia and axons to cross the site of injury to produce large swim. (**A-C**) *Left panels:* Total swim distance of control (black) vs transected larvae (red) generated from large movements (swim speed: ≥4 mm/s). Mean ± SD are shown. A two-way ANOVA using Tukey’s multiple comparison test was used to determine significance. *Right panels:* Quantification of swim distance from small movements classified by bridging across the injury site at the end of the experiment (5 dpi). An experimenter blinded to the swim analysis imaged larvae for the presence of a bridge. Mean ± SD is shown with individual fish plotted. Statistics for the observed phenotypes are in Table S3. (**D-F**) Total number of large movements of control (black) vs transected larvae (red) over the 10-min period of recording. Mean ± SD are shown. A two-way ANOVA using Tukey’s multiple comparison test was used to determine significance. 3 dpf: Control: n=28, SCI: n=28; 5 dpf: Control: n=29, SCI: n=30; 7 dpf: Control: 46, SCI: n=39 *p<0.05, **p<0.01, ***p<0.001, ****p<0.0001, ns: p≥0.4. Red text indicates comparisons between SCI 0 dpi and 5 dpi. Black text indicates comparisons between age- matched control and transected larvae.

**Figure 8.**
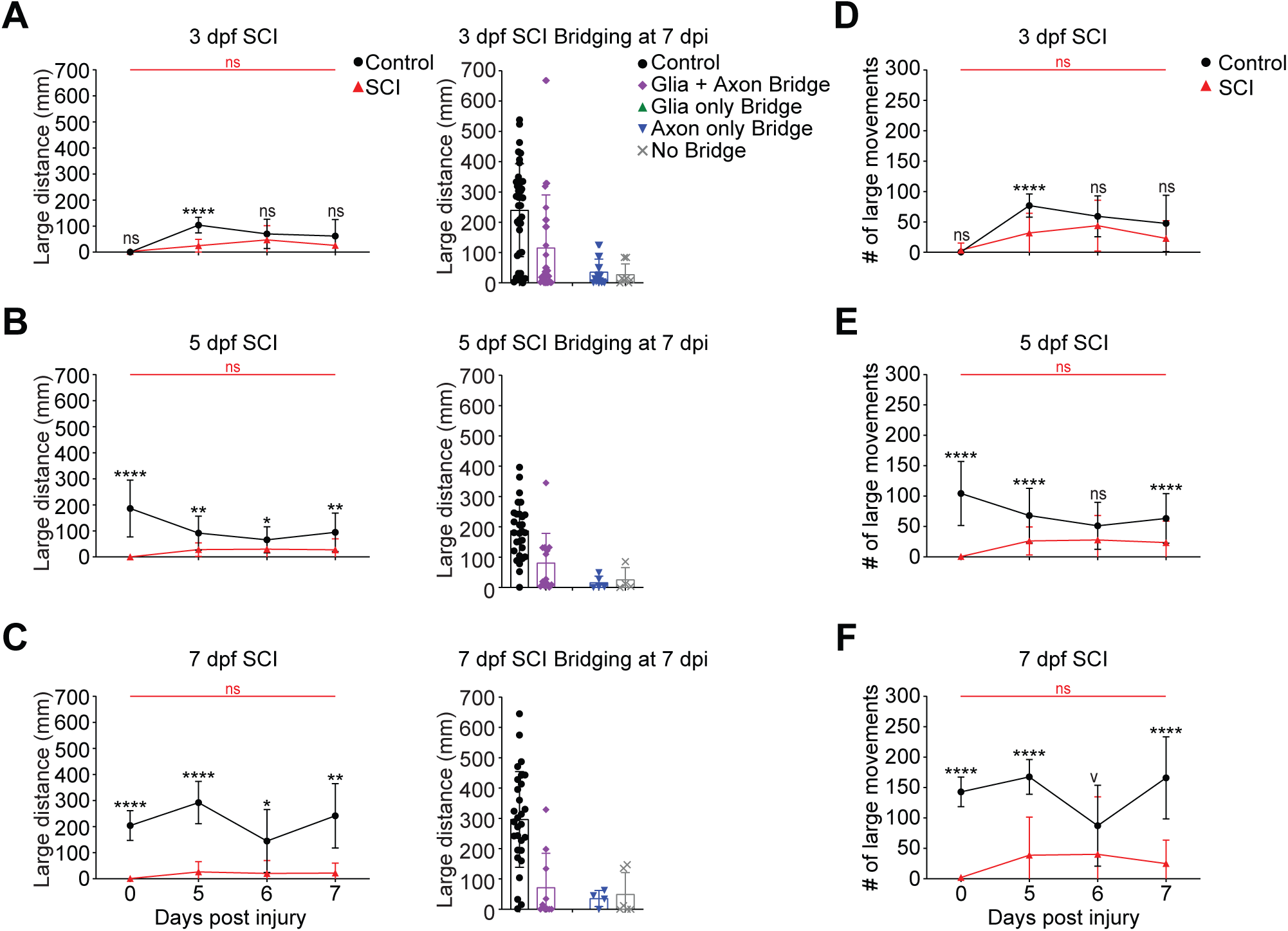
Additional recovery time increases number of larvae with glial and axonal bridging but did not affect swim distance. (**A-C**) *Left panels:* Total swim distance of control (black) vs transected larvae (red) generated from large movements (swim speed: ≥4 mm/s). Mean ± SD are shown. A two-way ANOVA using Tukey’s multiple comparison test was used to determine significance. *Right panels:* Quantification of swim distance from large movements classified by bridging across the injury site at the end of the experiment (5 dpi). An experimenter blinded to the swim analysis imaged larvae for the presence of a bridge. Mean ± SD is shown with individual fish plotted. Statistics for the observed phenotypes are in Table S4. (**D-F**) Total number of large movements of control (black) vs transected larvae (red) over the 10-min period of recording. Mean ± SD are shown. A two-way ANOVA using Tukey’s multiple comparison test was used to determine significance. 3 dpf: Control: n=47, SCI: n=59; 5 dpf: Control: n=29, SCI: n=22; 7 dpf: Control: 29, SCI: n=20 *p<0.05, **p<0.01, ***p<0.001, ****p<0.0001, ns: p≥0.1. Red text indicates comparisons between SCI 0 dpi and 5 dpi. Black text indicates comparisons between age-matched control and transected larvae.

## DISCUSSION

Motivated by the boom of SCI studies in various aged zebrafish, we systematically looked at SCI in three ages of larval zebrafish to identify similarities and differences of the cellular recovery process. Although the process of spinal cord regeneration is distinct from spinal development (Alper and Dorsky, 2022), we observed differences across the different ages of injury, indicating that there is an interplay between development and regeneration. For example, we found swim bladder development was impaired in larvae that were injured young (Fig. S1). This is perhaps unsurprising as both patterning of the spinal cord after injury (Reimer et al., 2009) and swim bladder development (Winata et al., 2009) require sonic hedgehog (Shh) signaling. However, few studies address whether and how resources are prioritized in developing animals across species to promote regeneration. Our results open the door to answering new experimental questions using this model organism, as larvae have many experimental advantages. For example, transgenic lines can be used to look at individual larvae such that a specific injury can be monitored from time of injury to regeneration instead of fixed samples. Factors such as the size of the initial injury and how much the injury area expands are challenging to assay in fixed tissues without a direct comparison to the initial injury.

SCI studies in zebrafish have focused on identifying the molecular drivers of axonal regeneration (Drake et al., 2023; Fang et al., 2014; Garcia et al., 2018; Ghosh and Hui, 2018; Keatinge et al., 2021; Li et al., 2020b; Noorimotlagh et al., 2017), circuit remodeling (Huang et al., 2022; Huang et al., 2021), glial bridging (Goldshmit et al., 2012; Klatt Shaw et al., 2021; Mokalled et al., 2016), and the immune response (Anguita-Salinas et al., 2019; de Sena-Tomas et al., 2024; Tsarouchas et al., 2018; Vandestadt et al., 2021). Our studies take a broad-strokes approach to look at the whole organism during the process of regeneration. This work opens the door to using larval zebrafish to study the mechanisms of known complications of SCI. For example, SCI is associated with cardiac dysfunction in mammals (Grigorean et al., 2009; Partida et al., 2016; Popa et al., 2010), including cardiac dysrhythmias, cardiac arrest, and hypotension. Our results indicate that larval zebrafish may exhibit some of these same dysfunctions (Fig. S1) providing a new model to address SCI complications. Although we did not look at other organ systems in this study, considering the anatomical similarities of the zebrafish to mammals, future studies looking at effects in the urinary (Kolvenbach et al., 2023; Outtandy et al., 2019) and gastrointestinal tract (Flores et al., 2020; Purifoy and Mruk, 2024; Sadler et al., 2013) are warranted.

We found differences in cytokine expression after transection in different ages. Activation of immune cells is one of the first responses after an injury. In larvae, neutrophils, macrophages, and microglia are all capable of secreting and responding to cytokines after SCI. Previous studies using 3 dpf larvae found an increase in the pro-inflammatory cytokine, *il-1*β within hours after injury. We observed the same for larvae transected at 3 dpf; however, *il-1*β wasn’t upregulated until four days after the injury in older larvae, indicating that the immune response may vary with maturity of the innate immune system. Given that both macrophages and microglia are still developing in the larval zebrafish (Xu et al., 2015; Zhao et al., 2024), it is perhaps unsurprising that older larvae exhibit a different cytokine profile after injury. Similarly, only larvae transected at 3 dpf and 5 dpf upregulated expression of TGF-β. Our results indicate that the immaturity of immune cells in younger larvae may account for the faster regeneration observed. One limitation of cytokine studies using qRT-PCR is that individual fish cannot be tracked throughout a study. Although we eliminated zebrafish from our study that had notochord damage or a cut width above 250 microns, we cannot guaranteed that our samples didn’t have different amounts of regeneration at each timepoint. Additional studies to discern the contribution of each cell type after SCI in older larvae and adults is warranted.

Whether axons require a glial bridge to cross the site of injury is still controversial (Cigliola et al., 2020). Axons severed during injury from neurons with cell bodies located in the brain, regrow and innervate their target to restore function (Becker et al., 1997); however, axonal regeneration is slowed in aging zebrafish (Graciarena et al., 2014). Ablation of GFAP+ cells with nitroreductase demonstrated that axons could cross the injury site in the absence of a glial bridge (Wehner et al., 2018) suggesting a glial bridge is not necessary for axonal regrowth; however, these studies were done on young larvae (3 dpf). Ablation of glial cells in adult zebrafish impairs glia bridging and axonal regrowth (Zhou et al., 2023). Although our studies do not try to answer this contested question, we did observe that a larger percentage of young larvae (3 dpf) were able to send axons across the site of injury independent of glial bridging than larvae transected at 5 dpf or 7 dpf. These differences again suggest that the developmental maturity of the larvae contributes to the differences reported in the literature. Therefore, studies looking at interactions between radial glia and axons at the injury border could shed light on whether the glial environment, metabolic state of radial glia, or other intrinsic properties dictate whether axons can cross the site of injury (Li et al., 2020a). Future studies combining transgenic lines with more efficient (Labbaf et al., 2022) and/or spatiotemporally controlled ablation (Mruk et al., 2020) could provide further insights into the axonal-glial interactions that occur during regrowth in larval animals. Lastly, for these studies we focused on recovery of free swim behavior. Consistent with the literature (Brustein et al., 2003; Drapeau et al., 2002), we found that 3 dpf larvae lacked coordinated movement independent of an injury. Manual transection of the spinal cord before coordinated movement is fully formed, may explain why 3 dpf larvae do not require glial or axons to cross the site of injury to recovery swim behavior. Our studies suggest that functional assays are more robust when larvae are transected after coordinated movement has developed.

Given that the number of laboratories studying SCI in zebrafish is steadily increasing, the need to standardize and expand these studies to include additional physiological considerations is vital for understanding the mechanisms of regeneration. We suggest that further investigation into cellular fate and molecular mechanisms in older larvae will improve the translational potential of this model with respect to CNS regeneration studies.

## METHODS

### Zebrafish husbandry

Adult zebrafish [wild-type AB and *Tg(gfap:EGFP)*], 3-18 months, were a generous gift from the Chen Lab (Stanford). Casper zebrafish were purchased from ZIRC (ZL1714). *Tg(olig2:dsRed)* were a generous gift from the Appel lab (UC Anschutz) (Kucenas et al., 2008). All zebrafish larvae were raised on a rotifer/brine shrimp diet starting at 5 days post fertilization (dpf) unless otherwise indicated (Purifoy and Mruk, 2024). Adults were maintained at 28.5°C on a 14:10 hour light:dark cycle and fed in the morning with Ziegler’s adult zebrafish diet and in the afternoon with brine shrimp. Embryos were staged as described previously (Kimmel et al., 1995). Embryos were obtained through natural matings and cultured at 28–30°C in E3 medium. To prevent pigmentation for live imaging experiments, the culture medium was also supplemented with 0.003% (w/v) N-phenylthiourea (PTU, Sigma-Aldrich #P7629). The IACUC committees at the University of Wyoming and East Carolina University approved all animal procedures.

### Line Generation

The generation of the Tol2-*elavl3:mCherry-CAAX* plasmid is previously described (Mruk et al., 2020). Transgenic zebrafish were generated using Tol2-mediated transgenesis (Abe et al., 2011; Suster et al., 2009). Plasmid DNA and Tol2 mRNA were premixed and co-injected into one-cell-stage embryos (50 pg of plasmid; 50 pg of mRNA). Fish were raised to adulthood and mated with Casper fish to identify founders with germline transmission, and those yielding F2 generations with monoallelic expression were used to establish the *Tg(elavl3:mCherry-CAAX*) transgenic line. The Tol2-*elavl3:mCherry-CAAX* plasmid and line are available upon request.

### Zebrafish transection

Larval zebrafish at 3, 5, or 7 dpf were mounted on flat slides in 2% low melting-point (LMP) agarose and briefly anesthetized using 0.01% tricaine (Western Chemical). The spinal cord was transected with a microblade (WPI#501731 or WPI#500249) at the level of the cloaca. Larvae were allowed to recover for approximately 2 hours in petri dishes filled with E2 buffer supplemented with penicillin/streptomycin (Pen/Strep, 100 unit, 100 ug/mL, Gibco #15140-122), before mounting and imaging (Briona and Dorsky, 2014b). Larvae were excluded if there was (1) an incomplete transection in the ventral spinal cord using the motor neuron tract *Tg(olig2:dsRed)* as a marker, (2) notochord damage during transection, or (3) larvae did not survive the length of the study. Larvae used for fluorescent imaging were also supplemented with 0.003% PTU. For multi-day experiments, E2 buffer was replaced daily. Larvae were not fed on the day of transection.

### Heartrate quantification

Live larvae were mounted in 1.5% LMP agarose on a glass slide and heartbeat was recorded via Olympus SZX16 stereoscope with an Olympus DP80 camera and SDF PLANO 1XPF objective based on ZebraPace (Gaur et al., 2018). Thirty second uncompressed AVI format videos were taken using Olympus cellSens software. Files were opened in FIJI software in grayscale mode without a virtual stack (Schindelin et al., 2012). A circular region of interest (ROI) was drawn on the AVI video file on the edge of the cardiac ventricle of the zebrafish larvae. Using ImageJ’s “plot-z-axis profile,” a time profile was generated via the pixel intensity change of the ROI. Peak detection and heartbeat counting was performed in MS Excel as previously described (Gaur et al., 2018). For each video, three 10-second sections were sampled and averaged to mediate random error. All videos were analyzed by a student blinded to the experimental conditions.

### Swim bladder inflation

Transgenic larvae from both *Tg(gfap:EGFP)* and *Tg(olig2:dsRed)* were transected as described. Larvae were observed each day, counting the number of animals with an inflated swim bladder each morning. Representative larvae from each condition were mounted in 1% LMP agarose and imaged on the Olympus SZX16 stereoscope with an Olympus DP80 camera and SDF PLANO 1XPF objective. Images were taken using the Olympus cellSens software.

### Imaging for Glial & Axonal Bridging

Heterozygous *Tg(elavl3:mCherry-CAAX)* larvae were crossed with *Tg(gfap:EGFP)* larvae to generate *Tg(elavl3:mCherry-CAAX; gfap:EGFP)* larvae. Larvae were transected as described above. Both transected and uncut controls were mounted in 1% LMP agarose and imaged on a Zeiss LSM 700 confocal microscope equipped with a 20x/NA 0.5 water-immersion objective. Z-stacks were generated from images taken at 2-5 µm intervals using the following settings: 2048x2048 pixels, 8 speed, 4 averaging. Maximum intensity projections were made in ImageJ and bridging was determined by observation of a complete track of fluorescence across the site of injury.

### Fluorescence intensity quantification

To quantify pixel intensity, raw green, fluorescent 20x z-stacks were opened in FIJI as maximum projections. The region of interest was selected using the segmented line selection tool at 200- point width. The Plot Profile analysis tool was then used to measure pixel intensity across the region of interest. Larvae with injuries larger than 250 micron, and larvae that did not survive the entire study were excluded from analysis.

### Acridine orange staining and cell counting

*Tg(olig2:dsRed)* embryos were transected as described. Live transected larvae were soaked in 5 µg/mL of acridine orange (Sigma) in E2 buffer in the dark for 1 hour at 28-29°C. Embryos were then washed 3 times for 5 minutes each in fresh E2 buffer, mounted on glass slides in 1% LMP agarose, and imaged on a Leica DM6D epifluorescence microscope with a Leica DFC900GT camera and CoolLED pE-300 Ultra light source or on a Zeiss LSM 700 confocal microscope equipped with a MA-PMT with 20x/NA 0.5 water immersion objective. Z-stacks were generated from images taken at 2-5 µm intervals using the following settings: 1024x1024 pixels, 9 speed, 4 averaging. Total number of AO+ cells in the spinal tract were counted by two individual researchers and the average of the two researchers for each larvae plotted.

### Quantitative PCR

*Tg(gfap:EGFP)* larvae were transected as described. Five larvae for each condition for each time point were selected and cDNA generated as described above. cDNA samples were cleaned using ethanol precipitation. Approximately 200 ng of each cDNA sample was used as templates for qPCR using the Maxima SYBR Green/ROX master mix (Thermo Scientific #K0221). Target sequences for qPCR were amplified using primers for *gfap*, *ctgfa, gapdh*, *tgf-*β*1*, *tgf-*β*3*, *tnf-*α, and *il-1*β (Table 2).(DiMuccio et al., 2005) (Tsarouchas et al., 2018). Relative mRNA levels were determined using the ΔΔC_t_ method. The relative fold change was calculated using 2^-ΔΔCt^.

**Table 2.**
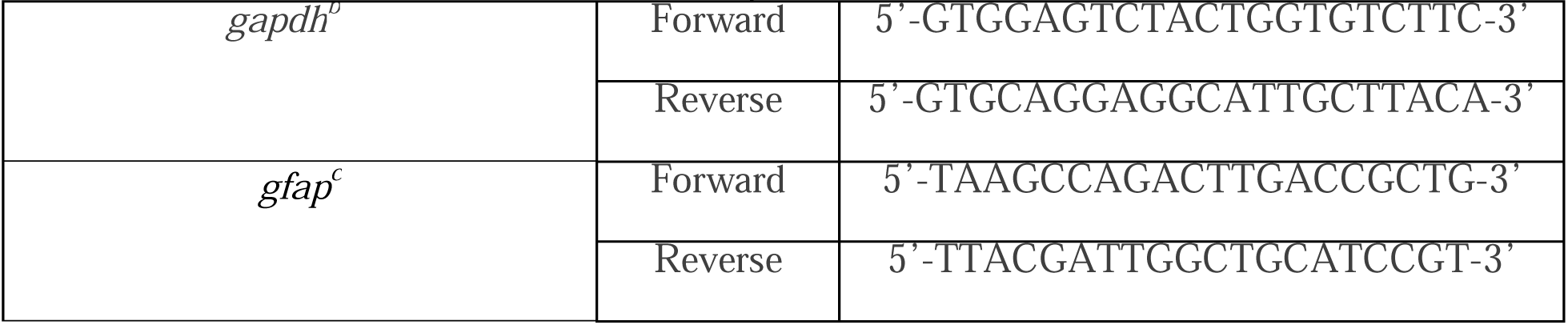
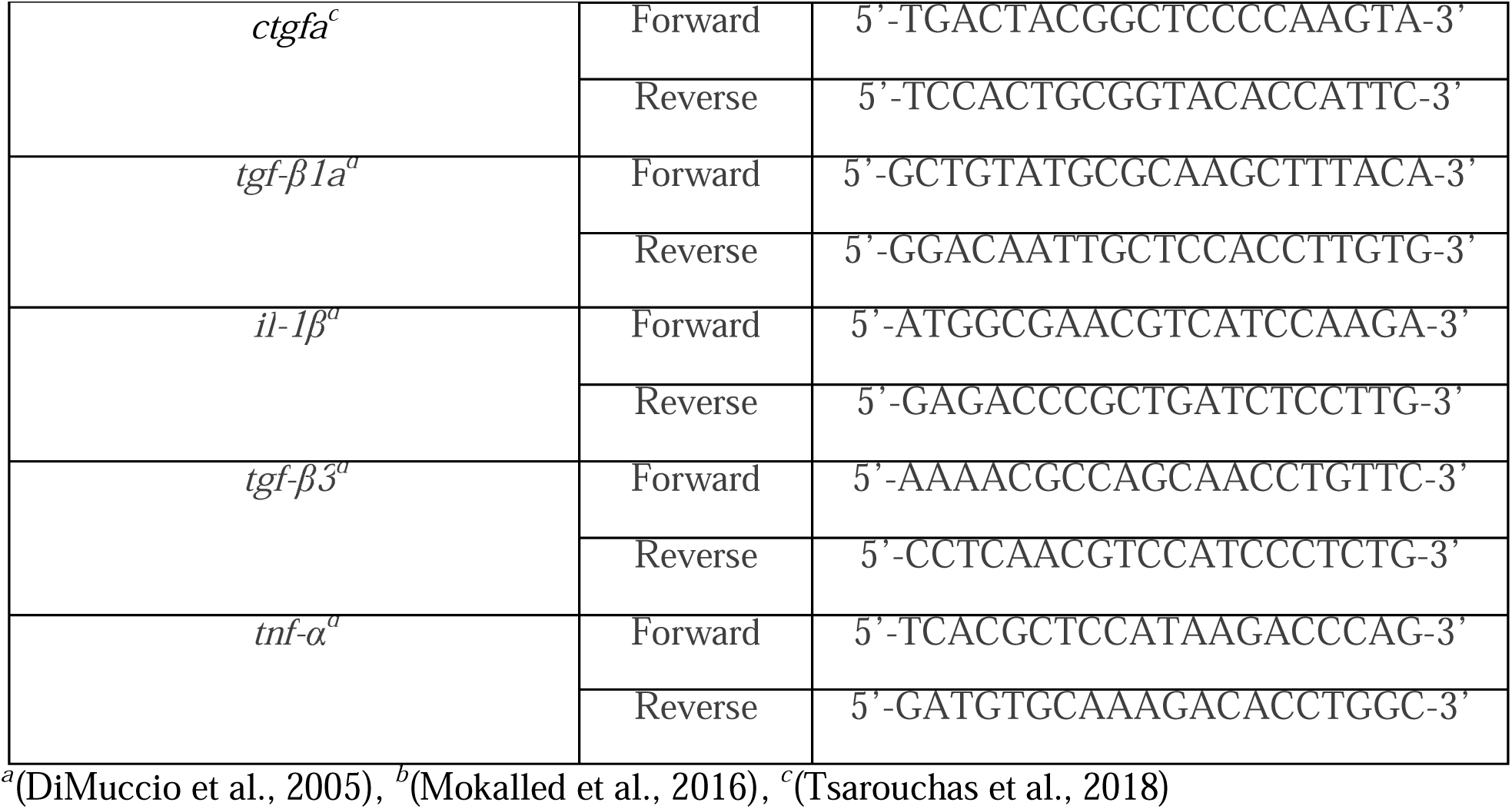
qRT-PCR primers used in this study.

### Locomotion behavior

*Tg(gfap:EGFP; elav3:mCherry-CAAX)* larvae were transected as described and locomotion of individual fish recorded through 14 dpf. Larval zebrafish reliably show activity on transition to a dark environment from a lit environment and are more active in the morning beginning around 10 am and reach their lowest activity in the evening by 5 pm (MacPhail et al., 2009). Behavior experiments were conducted 10:00 am–noon EST for all ages. A clear 48-well plate was used with one fish per well in a fixed volume of 1 mL of E2 supplemented with Pen/Strep for the locomotion assay. Locomotion was assessed using a Zebrabox and its ViewPoint LS tracking software (Viewpoint Life Sciences, Lyon, France). Larvae were acclimated to the test environment for 10 minutes, followed by a 10-minute session in a dark environment. Locomotion speed from various movement types (twitch: 0-2 mm/s; small: 2-4 mm/s; large: >4 mm/s) was video recorded. Following behavior experiments, larvae were imaged and scored by an independent experimenter using Leica DM6D epifluorescence microscope with a Leica K8 camera and CoolLED pE-300 Ultra light source.

### Statistics

For all zebrafish experiments, at least two breeding tanks, each containing 2 to 4 males and 2 to 4 females from separate stocks, were set up to generate embryos. Embryos from each tank were randomly distributed across tested conditions, and unfertilized and developmentally abnormal embryos were removed prior to transection or compound treatment. Samples sizes were calculated for a 30% effect with 80% power. For all graphs, values for individual fish are plotted, and data is presented as the mean ± standard deviation (SD). For statistical testing, each distribution was assessed using the Shapiro-Wilk test and determined to be non-normal. Significant differences were determined using either an unpaired Mann-Whitney t-test or a Kruskal-Wallis ANOVA test with a post hoc Dunn’s test. For qPCR technical replicates outliers were determined using the Grubb’s test with α = 0.05 and significant differences were determined using a two-way ANOVA test with Tukey’s multiple comparisons test. P values of < 0.05 were considered statistically significant. Graphs were generated using GraphPad Prism 9 (Dotmatics) software. Additional experimental statistics for the phenotypic distributions are reported in Tables S1-S4.

## Supporting information

Supplemental Packet

## ACKNOWLEDGMENTS

We gratefully acknowledge financial support from NIH P20GM121310 and R03NS136719 (K.M.) and the McNair Scholars program (S.M.L). Confocal imaging was supported in part by the University of Wyoming’s Integrated Microscopy Core (NIH P20GM121310).

## AUTHOR CONTRIBUTIONS

Conceptualization: K.L.U, K.M.; Methodology: W.J.W., K.L.U., P.I.G., K.B.L., S.M.L., K.M.; Validation: W.J.W., K.L.U., P.I.G., K.B.L., K.M.; Formal analysis: W.J.W., K.L.U., P.I.G., K.B.L, S.M.L., T.P.R., C.S. K.M.; Investigation: W.J.W., K.L.U., P.I.G., K.B.L, K.M.; Resources: W.J.W., T.P.R. Writing – original draft: W.J.W, K.L.U, P.I.G., K.B.L., K.M.; Writing - review & editing: W.J.W., K.L.U, P.I.G., K.B.L, S.M.L., T.P.R., C.S. K.M.; Visualization: W.J.W., K.L.U., P.I.G., K.B.L., S.M.L., K.M. Supervision: K.M.; Project administration: K.M.; Funding acquisition: K.M.

## SUPPORTING INFORMATION

Tables S1-S3 Figs. S1-S3

## REFERENCES

1. Abe, G., Suster, M.L., Kawakami, K., 2011. Tol2-mediated transgenesis, gene trapping, enhancer trapping, and the Gal4-UAS system. Methods in cell biology 104, 23–49.

2. Alper, S.R., Dorsky, R.I., 2022. Unique advantages of zebrafish larvae as a model for spinal cord regeneration. Frontiers in molecular neuroscience 15, 983336.

3. Anguita-Salinas, C., Sánchez, M., Morales, R.A., Ceci, M.L., Rojas-Benítez, D., Allende, M.L., 2019. Cellular Dynamics during Spinal Cord Regeneration in Larval Zebrafish. Developmental neuroscience 41, 112–122.

4. Becker, T., Wullimann, M.F., Becker, C.G., Bernhardt, R.R., Schachner, M., 1997. Axonal regrowth after spinal cord transection in adult zebrafish. The Journal of comparative neurology 377, 577–595.

5. Bensimon-Brito, A., Cardeira, J., Cancela, M.L., Huysseune, A., Witten, P.E., 2012. Distinct patterns of notochord mineralization in zebrafish coincide with the localization of Osteocalcin isoform 1 during early vertebral centra formation. BMC developmental biology 12, 28.

6. Berg, E.M., Mrowka, L., Bertuzzi, M., Madrid, D., Picton, L.D., El Manira, A., 2023. Brainstem circuits encoding start, speed, and duration of swimming in adult zebrafish. Neuron 111, 372–386 e374.

7. Bin, J.M., Lyons, D.A., 2016. Imaging Myelination In Vivo Using Transparent Animal Models. Brain plasticity (Amsterdam, Netherlands) 2, 3–29.

8. Briona, L.K., Dorsky, R.I., 2014a. Radial glial progenitors repair the zebrafish spinal cord following transection. Experimental neurology 256, 81–92.

9. Briona, L.K., Dorsky, R.I., 2014b. Spinal cord transection in the larval zebrafish. Journal of visualized experiments : JoVE.

10. Brösamle, C., Halpern, M.E., 2002. Characterization of myelination in the developing zebrafish. Glia 39, 47–57.

11. Brown, D.R., Samsa, L.A., Qian, L., Liu, J., 2016. Advances in the Study of Heart Development and Disease Using Zebrafish. Journal of cardiovascular development and disease 3.

12. Brustein, E., Saint-Amant, L., Buss, R.R., Chong, M., McDearmid, J.R., Drapeau, P., 2003. Steps during the development of the zebrafish locomotor network. Journal of physiology, Paris 97, 77–86.

13. Cavone, L., McCann, T., Drake, L.K., Aguzzi, E.A., Oprişoreanu, A.M., Pedersen, E., Sandi, S., Selvarajah, J., Tsarouchas, T.M., Wehner, D., Keatinge, M., Mysiak, K.S., Henderson, B.E.P., Dobie, R., Henderson, N.C., Becker, T., Becker, C.G., 2021. A unique macrophage subpopulation signals directly to progenitor cells to promote regenerative neurogenesis in the zebrafish spinal cord. Developmental cell 56, 1617–1630 e1616.

14. Cigliola, V., Becker, C.J., Poss, K.D., 2020. Building bridges, not walls: spinal cord regeneration in zebrafish. Disease models & mechanisms 13.

15. de Sena-Tomas, C., Rebola Lameira, L., Rebocho da Costa, M., Naique Taborda, P., Laborde, A., Orger, M., de Oliveira, S., Saude, L., 2024. Neutrophil immune profile guides spinal cord regeneration in zebrafish. Brain Behav Immun 120, 514–531.

16. DiMuccio, T., Mukai, S.T., Clelland, E., Kohli, G., Cuartero, M., Wu, T., Peng, C., 2005. Cloning of a second form of activin-betaA cDNA and regulation of activin-betaA subunits and activin type II receptor mRNA expression by gonadotropin in the zebrafish ovary. General and comparative endocrinology 143, 287–299.

17. Drake, L.K., Keatinge, M., Tsarouchas, T.M., Becker, C.G., Lyons, D.A., Becker, T., 2023. Rapid Testing of Gene Function in Axonal Regeneration After Spinal Cord Injury Using Larval Zebrafish. Methods in molecular biology (Clifton, N.J.) 2636, 263–277.

18. Drapeau, P., Saint-Amant, L., Buss, R.R., Chong, M., McDearmid, J.R., Brustein, E., 2002. Development of the locomotor network in zebrafish. Progress in neurobiology 68, 85–111.

19. Fang, P., Lin, J.F., Pan, H.C., Shen, Y.Q., Schachner, M., 2012. A surgery protocol for adult zebrafish spinal cord injury. Journal of genetics and genomics = Yi chuan xue bao 39, 481–487.

20. Fang, P., Pan, H.C., Lin, S.L., Zhang, W.Q., Rauvala, H., Schachner, M., Shen, Y.Q., 2014. HMGB1 contributes to regeneration after spinal cord injury in adult zebrafish. Molecular neurobiology 49, 472–483.

21. Flores, E.M., Nguyen, A.T., Odem, M.A., Eisenhoffer, G.T., Krachler, A.M., 2020. The zebrafish as a model for gastrointestinal tract-microbe interactions. Cellular microbiology 22, e13152.

22. Garcia, A.L., Udeh, A., Kalahasty, K., Hackam, A.S., 2018. A growing field: The regulation of axonal regeneration by Wnt signaling. Neural regeneration research 13, 43–52.

23. Gaur, H., Pullaguri, N., Nema, S., Purushothaman, S., Bhargava, Y., Bhargava, A., 2018. ZebraPace: An Open-Source Method for Cardiac-Rhythm Estimation in Untethered Zebrafish Larvae. Zebrafish 15, 254–262.

24. Ghosh, S., Hui, S.P., 2018. Axonal regeneration in zebrafish spinal cord. Regeneration (Oxford, England) 5, 43–60.

25. Goldshmit, Y., Sztal, T.E., Jusuf, P.R., Hall, T.E., Nguyen-Chi, M., Currie, P.D., 2012. Fgf-dependent glial cell bridges facilitate spinal cord regeneration in zebrafish. The Journal of neuroscience : the official journal of the Society for Neuroscience 32, 7477–7492.

26. Gollmann-Tepeköylü, C., Nägele, F., Graber, M., Pölzl, L., Lobenwein, D., Hirsch, J., An, A., Irschick, R., Röhrs, B., Kremser, C., Hackl, H., Huber, R., Venezia, S., Hercher, D., Fritsch, H., Bonaros, N., Stefanova, N., Tancevski, I., Meyer, D., Grimm, M., Holfeld, J., 2020. Shock waves promote spinal cord repair via TLR3. JCI insight 5.

27. Graciarena, M., Dambly-Chaudière, C., Ghysen, A., 2014. Dynamics of axonal regeneration in adult and aging zebrafish reveal the promoting effect of a first lesion. Proceedings of the National Academy of Sciences of the United States of America 111, 1610–1615.

28. Grigorean, V.T., Sandu, A.M., Popescu, M., Iacobini, M.A., Stoian, R., Neascu, C., Strambu, V., Popa, F., 2009. Cardiac dysfunctions following spinal cord injury. Journal of medicine and life 2, 133–145.

29. Hernandez, R.E., Galitan, L., Cameron, J., Goodwin, N., Ramakrishnan, L., 2018. Delay of Initial Feeding of Zebrafish Larvae Until 8 Days Postfertilization Has No Impact on Survival or Growth Through the Juvenile Stage. Zebrafish 15, 515–518.

30. Hossainian, D., Shao, E., Jiao, B., Ilin, V.A., Parris, R.S., Zhou, Y., Bai, Q., Burton, E.A., 2022. Quantification of functional recovery in a larval zebrafish model of spinal cord injury. Journal of neuroscience research 100, 2044–2054.

31. Huang, C.X., Wang, Z., Cheng, J., Zhu, Z., Guan, N.N., Song, J., 2022. De novo establishment of circuit modules restores locomotion after spinal cord injury in adult zebrafish. Cell reports 41, 111535.

32. Huang, C.X., Zhao, Y., Mao, J., Wang, Z., Xu, L., Cheng, J., Guan, N.N., Song, J., 2021. An injury-induced serotonergic neuron subpopulation contributes to axon regrowth and function restoration after spinal cord injury in zebrafish. Nature communications 12, 7093.

33. Hui, S.P., Dutta, A., Ghosh, S., 2010. Cellular response after crush injury in adult zebrafish spinal cord. Developmental dynamics : an official publication of the American Association of Anatomists 239, 2962–2979.

34. Hui, S.P., Sengupta, D., Lee, S.G., Sen, T., Kundu, S., Mathavan, S., Ghosh, S., 2014. Genome wide expression profiling during spinal cord regeneration identifies comprehensive cellular responses in zebrafish. PloS one 9, e84212.

35. Keatinge, M., Tsarouchas, T.M., Munir, T., Porter, N.J., Larraz, J., Gianni, D., Tsai, H.H., Becker, C.G., Lyons, D.A., Becker, T., 2021. CRISPR gRNA phenotypic screening in zebrafish reveals pro-regenerative genes in spinal cord injury. PLoS genetics 17, e1009515.

36. Kim, H., Kim, S., Chung, A.Y., Bae, Y.K., Hibi, M., Lim, C.S., Park, H.C., 2008. Notch-regulated perineurium development from zebrafish spinal cord. Neuroscience letters 448, 240–244.

37. Kimmel, C.B., Ballard, W.W., Kimmel, S.R., Ullmann, B., Schilling, T.F., 1995. Stages of embryonic development of the zebrafish. Developmental dynamics : an official publication of the American Association of Anatomists 203, 253–310.

38. Kirby, B.B., Takada, N., Latimer, A.J., Shin, J., Carney, T.J., Kelsh, R.N., Appel, B., 2006. In vivo time-lapse imaging shows dynamic oligodendrocyte progenitor behavior during zebrafish development. Nature neuroscience 9, 1506–1511.

39. Kishore, S., Bagnall, M.W., McLean, D.L., 2014. Systematic shifts in the balance of excitation and inhibition coordinate the activity of axial motor pools at different speeds of locomotion. The Journal of neuroscience : the official journal of the Society for Neuroscience 34, 14046–14054.

40. Klatt Shaw, D., Saraswathy, V.M., Zhou, L., McAdow, A.R., Burris, B., Butka, E., Morris, S.A., Dietmann, S., Mokalled, M.H., 2021. Localized EMT reprograms glial progenitors to promote spinal cord repair. Developmental cell 56, 613–626 e617.

41. Kolvenbach, C.M., Dworschak, G.C., Rieke, J.M., Woolf, A.S., Reutter, H., Odermatt, B., Hilger, A.C., 2023. Modelling human lower urinary tract malformations in zebrafish. Molecular and cellular pediatrics 10, 2.

42. Kucenas, S., Takada, N., Park, H.C., Woodruff, E., Broadie, K., Appel, B., 2008. CNS-derived glia ensheath peripheral nerves and mediate motor root development. Nature neuroscience 11, 143–151.

43. Labbaf, Z., Petratou, K., Ermlich, L., Backer, W., Tarbashevich, K., Reichman-Fried, M., Luschnig, S., Schulte-Merker, S., Raz, E., 2022. A robust and tunable system for targeted cell ablation in developing embryos. Developmental cell 57, 2026–2040 e2025.

44. Li, F., Sami, A., Noristani, H.N., Slattery, K., Qiu, J., Groves, T., Wang, S., Veerasammy, K., Chen, Y.X., Morales, J., Haynes, P., Sehgal, A., He, Y., Li, S., Song, Y., 2020a. Glial Metabolic Rewiring Promotes Axon Regeneration and Functional Recovery in the Central Nervous System. Cell metabolism 32, 767–785 e767.

45. Li, J.H., Shi, Z.J., Li, Y., Pan, B., Yuan, S.Y., Shi, L.L., Hao, Y., Cao, F.J., Feng, S.Q., 2020b. Bioinformatic identification of key candidate genes and pathways in axon regeneration after spinal cord injury in zebrafish. Neural regeneration research 15, 103–111.

46. MacPhail, R.C., Brooks, J., Hunter, D.L., Padnos, B., Irons, T.D., Padilla, S., 2009. Locomotion in larval zebrafish: Influence of time of day, lighting and ethanol. Neurotoxicology 30, 52–58.

47. Matsuoka, R.L., Marass, M., Avdesh, A., Helker, C.S., Maischein, H.M., Grosse, A.S., Kaur, H., Lawson, N.D., Herzog, W., Stainier, D.Y., 2016. Radial glia regulate vascular patterning around the developing spinal cord. eLife 5.

48. Mizuno, Y., Mochizuki, H., Sugita, Y., Goto, K., 1998. Apoptosis in neurodegenerative disorders. Internal medicine (Tokyo, Japan) 37, 192–193.

49. Mokalled, M.H., Patra, C., Dickson, A.L., Endo, T., Stainier, D.Y., Poss, K.D., 2016. Injury-induced ctgfa directs glial bridging and spinal cord regeneration in zebrafish. Science (New York, N.Y.) 354, 630–634.

50. Mruk, K., Ciepla, P., Piza, P.A., Alnaqib, M.A., Chen, J.K., 2020. Targeted cell ablation in zebrafish using optogenetic transcriptional control. Development (Cambridge, England) 147.

51. Noorimotlagh, Z., Babaie, M., Safdarian, M., Ghadiri, T., Rahimi-Movaghar, V., 2017. Mechanisms of spinal cord injury regeneration in zebrafish: a systematic review. Iranian journal of basic medical sciences 20, 1287–1296.

52. Ohnmacht, J., Yang, Y., Maurer, G.W., Barreiro-Iglesias, A., Tsarouchas, T.M., Wehner, D., Sieger, D., Becker, C.G., Becker, T., 2016. Spinal motor neurons are regenerated after mechanical lesion and genetic ablation in larval zebrafish. Development (Cambridge, England) 143, 1464–1474.

53. Outtandy, P., Russell, C., Kleta, R., Bockenhauer, D., 2019. Zebrafish as a model for kidney function and disease. Pediatric nephrology (Berlin, Germany) 34, 751–762.

54. Panula, P., Sallinen, V., Sundvik, M., Kolehmainen, J., Torkko, V., Tiittula, A., Moshnyakov, M., Podlasz, P., 2006. Modulatory neurotransmitter systems and behavior: towards zebrafish models of neurodegenerative diseases. Zebrafish 3, 235–247.

55. Parichy, D.M., Elizondo, M.R., Mills, M.G., Gordon, T.N., Engeszer, R.E., 2009. Normal table of postembryonic zebrafish development: staging by externally visible anatomy of the living fish. Developmental dynamics : an official publication of the American Association of Anatomists 238, 2975–3015.

56. Park, H.C., Mehta, A., Richardson, J.S., Appel, B., 2002. olig2 is required for zebrafish primary motor neuron and oligodendrocyte development. Developmental biology 248, 356–368.

57. Partida, E., Mironets, E., Hou, S., Tom, V.J., 2016. Cardiovascular dysfunction following spinal cord injury. Neural regeneration research 11, 189–194.

58. Popa, C., Popa, F., Grigorean, V.T., Onose, G., Sandu, A.M., Popescu, M., Burnei, G., Strambu, V., Sinescu, C., 2010. Vascular dysfunctions following spinal cord injury. Journal of medicine and life 3, 275–285.

59. Preston, M.A., Macklin, W.B., 2015. Zebrafish as a model to investigate CNS myelination. Glia 63, 177–193.

60. Purifoy, E.J., Mruk, K., 2024. Differential Roles of Diet on Development and Spinal Cord Regeneration in Larval Zebrafish. Zebrafish 21, 214–222.

61. Reimer, M.M., Kuscha, V., Wyatt, C., Sörensen, I., Frank, R.E., Knüwer, M., Becker, T., Becker, C.G., 2009. Sonic hedgehog is a polarized signal for motor neuron regeneration in adult zebrafish. The Journal of neuroscience : the official journal of the Society for Neuroscience 29, 15073–15082.

62. Reimer, M.M., Norris, A., Ohnmacht, J., Patani, R., Zhong, Z., Dias, T.B., Kuscha, V., Scott, A.L., Chen, Y.C., Rozov, S., Frazer, S.L., Wyatt, C., Higashijima, S., Patton, E.E., Panula, P., Chandran, S., Becker, T., Becker, C.G., 2013. Dopamine from the brain promotes spinal motor neuron generation during development and adult regeneration. Developmental cell 25, 478–491.

63. Sadler, K.C., Rawls, J.F., Farber, S.A., 2013. Getting the inside tract: new frontiers in zebrafish digestive system biology. Zebrafish 10, 129–131.

64. Schindelin, J., Arganda-Carreras, I., Frise, E., Kaynig, V., Longair, M., Pietzsch, T., Preibisch, S., Rueden, C., Saalfeld, S., Schmid, B., Tinevez, J.Y., White, D.J., Hartenstein, V., Eliceiri, K., Tomancak, P., Cardona, A., 2012. Fiji: an open-source platform for biological-image analysis. Nature methods 9, 676-682.

65. Severi, K.E., Portugues, R., Marques, J.C., O’Malley, D.M., Orger, M.B., Engert, F., 2014. Neural control and modulation of swimming speed in the larval zebrafish. Neuron 83, 692–707.

66. Suster, M.L., Kikuta, H., Urasaki, A., Asakawa, K., Kawakami, K., 2009. Transgenesis in zebrafish with the tol2 transposon system. Methods in molecular biology (Clifton, N.J.) 561, 41–63.

67. Tsarouchas, T.M., Wehner, D., Cavone, L., Munir, T., Keatinge, M., Lambertus, M., Underhill, A., Barrett, T., Kassapis, E., Ogryzko, N., Feng, Y., van Ham, T.J., Becker, T., Becker, C.G., 2018. Dynamic control of proinflammatory cytokines Il-1β and Tnf-α by macrophages in zebrafish spinal cord regeneration. Nature communications 9, 4670.

68. Vajn, K., Suler, D., Plunkett, J.A., Oudega, M., 2014. Temporal profile of endogenous anatomical repair and functional recovery following spinal cord injury in adult zebrafish. PloS one 9, e105857.

69. van Raamsdonk, W., Smit-Onel, M.J., Maslam, S., Velzing, E., de Heus, R., 1998. Changes in the synaptology of spinal motoneurons in zebrafish following spinal cord transection. Acta histochemica 100, 133–148.

70. Vandestadt, C., Vanwalleghem, G.C., Khabooshan, M.A., Douek, A.M., Castillo, H.A., Li, M., Schulze, K., Don, E., Stamatis, S.A., Ratnadiwakara, M., Änkö, M.L., Scott, E.K., Kaslin, J., 2021. RNA-induced inflammation and migration of precursor neurons initiates neuronal circuit regeneration in zebrafish. Developmental cell 56, 2364–2380 e2368.

71. Wehner, D., Becker, T., Becker, C.G., 2018. Restoration of anatomical continuity after spinal cord transection depends on Wnt/β-catenin signaling in larval zebrafish. Data in brief 16, 65–70.

72. Winata, C.L., Korzh, S., Kondrychyn, I., Zheng, W., Korzh, V., Gong, Z., 2009. Development of zebrafish swimbladder: The requirement of Hedgehog signaling in specification and organization of the three tissue layers. Developmental biology 331, 222–236.

73. Xu, J., Zhu, L., He, S., Wu, Y., Jin, W., Yu, T., Qu, J.Y., Wen, Z., 2015. Temporal-Spatial Resolution Fate Mapping Reveals Distinct Origins for Embryonic and Adult Microglia in Zebrafish. Developmental cell 34, 632–641.

74. Zhao, C., Yang, Z., Li, Y., Wen, Z., 2024. Macrophages in tissue repair and regeneration: insights from zebrafish. Cell Regen 13, 12.

75. Zhou, L., McAdow, A.R., Yamada, H., Burris, B., Klatt Shaw, D., Oonk, K., Poss, K.D., Mokalled, M.H., 2023. Progenitor-derived glia are required for spinal cord regeneration in zebrafish. Development (Cambridge, England) 150.

