## Supplemental Packet for "Effects of age on the response to spinal cord injury: optimizing the larval zebrafish model"

**Table S1. Experimental statistics for number of apoptotic cells in zebrafish larvae** (related to Figure 1).

| Panel | Transgenic Line (heterozygous) | Stage | Treatment | Total number of animals |
|---|---|---|---|---|
| E | *Tg(olig2:dsRed)* | 3 dpf 12 hpi (4 dpf) | Control | 24 |
| E | *Tg(olig2:dsRed)* | | SCI | 27 |
| E | *Tg(olig2:dsRed)* | 3 dpf 24 hpi (4 dpf) | Control | 24 |
| E | *Tg(olig2:dsRed)* | | SCI | 22 |
| E | *Tg(olig2:dsRed)* | 3 dpf 48 hpi (5 dpf) | Control | 26 |
| E | *Tg(olig2:dsRed)* | | SCI | 22 |
| E | *Tg(olig2:dsRed)* | 3 dpf 72 hpi (6 dpf) | Control | 24 |
| E | *Tg(olig2:dsRed)* | | SCI | 21 |
| E | *Tg(olig2:dsRed)* | 3 dpf 96 hpi (7 dpf) | Control | 25 |
| E | *Tg(olig2:dsRed)* | | SCI | 21 |
| E | *Tg(olig2:dsRed)* | 3 dpf 120 hpi (8 dpf) | Control | 23 |
| E | *Tg(olig2:dsRed)* | | SCI | 24 |
| E | *Tg(olig2:dsRed)* | 5 dpf 12 hpi (6 dpf) | Control | 22 |
| E | *Tg(olig2:dsRed)* | | SCI | 24 |
| E | *Tg(olig2:dsRed)* | 5 dpf 24 hpi (6 dpf) | Control | 23 |
| E | *Tg(olig2:dsRed)* | | SCI | 33 |
| E | *Tg(olig2:dsRed)* | 5 dpf 48 hpi (7 dpf) | Control | 25 |
| E | *Tg(olig2:dsRed)* | | SCI | 25 |
| E | *Tg(olig2:dsRed)* | 5 dpf 72 hpi (8 dpf) | Control | 23 |
| E | *Tg(olig2:dsRed)* | | SCI | 21 |
| E | *Tg(olig2:dsRed)* | 5 dpf 96 hpi (9 dpf) | Control | 23 |
| E | *Tg(olig2:dsRed)* | | SCI | 21 |
| E | *Tg(olig2:dsRed)* | 5 dpf 120 hpi (10 dpf) | Control | 23 |
| E | *Tg(olig2:dsRed)* | | SCI | 26 |
| E | *Tg(olig2:dsRed)* | 7 dpf 12 hpi (8 dpf) | Control | 29 |
| E | *Tg(olig2:dsRed)* | | SCI | 33 |
| E | *Tg(olig2:dsRed)* | 7 dpf 24 hpi (8 dpf) | Control | 28 |
| E | *Tg(olig2:dsRed)* | | SCI | 24 |
| E | *Tg(olig2:dsRed)* | 7 dpf 48 hpi (9 dpf) | Control | 24 |
| E | *Tg(olig2:dsRed)* | | SCI | 35 |
| E | *Tg(olig2:dsRed)* | 7 dpf 72 hpi (10 dpf) | Control | 32 |
| E | *Tg(olig2:dsRed)* | | SCI | 24 |
| E | *Tg(olig2:dsRed)* | 7 dpf 96 hpi (11 dpf) | Control | 26 |
| E | *Tg(olig2:dsRed)* | | SCI | 30 |
| E | *Tg(olig2:dsRed)* | 7 dpf 120 hpi (12 dpf) | Control | 27 |
| E | *Tg(olig2:dsRed)* | | SCI | 35 |

**Table S2. Experimental statistics for glial and axonal bridging** (related to Figure 3).

| Panel | Transgenic Line (heterozygous) | Stage | Treatment – Condition | Total number of animals |
|---|---|---|---|---|
| B | *Tg(gfap:EGFP; elavl3:mCherry-CAAX)* | 3 dpf 0 dpi (3 dpf) | SCI – no bridging | 18 |
| B | *Tg(gfap:EGFP; elavl3:mCherry-CAAX)* | 3 dpf 1 dpi (4 dpf) | SCI – no bridging | 16 |
| B | *Tg(gfap:EGFP; elavl3:mCherry-CAAX)* | 3 dpf 1 dpi (4 dpf) | SCI – axonal bridging only | 1 |
| B | *Tg(gfap:EGFP; elavl3:mCherry-CAAX)* | 3 dpf 1 dpi (4 dpf) | SCI – glial & axonal bridging | 1 |
| B | *Tg(gfap:EGFP; elavl3:mCherry-CAAX)* | 3 dpf 2 dpi (5 dpf) | SCI – no bridging | 8 |
| B | *Tg(gfap:EGFP; elavl3:mCherry-CAAX)* | 3 dpf 2 dpi (5 dpf) | SCI – axonal bridging only | 0 |
| B | *Tg(gfap:EGFP; elavl3:mCherry-CAAX)* | 3 dpf 2 dpi (5 dpf) | SCI – glial & axonal bridging | 10 |
| B | *Tg(gfap:EGFP; elavl3:mCherry-CAAX)* | 3 dpf 3 dpi (6 dpf) | SCI – no bridging | 3 |
| B | *Tg(gfap:EGFP; elavl3:mCherry-CAAX)* | 3 dpf 3 dpi (6 dpf) | SCI – axonal bridging only | 4 |
| B | *Tg(gfap:EGFP; elavl3:mCherry-CAAX)* | 3 dpf 3 dpi (6 dpf) | SCI – glial & axonal bridging | 11 |
| B | *Tg(gfap:EGFP; elavl3:mCherry-CAAX)* | 3 dpf 4 dpi (7 dpf) | SCI – no bridging | 2 |
| B | *Tg(gfap:EGFP; elavl3:mCherry-CAAX)* | 3 dpf 4 dpi (7 dpf) | SCI – axonal bridging only | 4 |
| B | *Tg(gfap:EGFP; elavl3:mCherry-CAAX)* | 3 dpf 4 dpi (7 dpf) | SCI – glial & axonal bridging | 12 |
| B | *Tg(gfap:EGFP; elavl3:mCherry-CAAX)* | 3 dpf 5 dpi (8 dpf) | SCI – no bridging | 2 |
| B | *Tg(gfap:EGFP; elavl3:mCherry-CAAX)* | 3 dpf 5 dpi (8 dpf) | SCI – axonal bridging only | 4 |
| B | *Tg(gfap:EGFP; elavl3:mCherry-CAAX)* | 3 dpf 5 dpi (8 dpf) | SCI – glial & axonal bridging | 12 |
| B | *Tg(gfap:EGFP; elavl3:mCherry-CAAX)* | 5 dpf 0 dpi (5 dpf) | SCI – no bridging | 20 |
| B | *Tg(gfap:EGFP; elavl3:mCherry-CAAX)* | 5 dpf 1 dpi (6 dpf) | SCI – no bridging | 20 |
| B | *Tg(gfap:EGFP; elavl3:mCherry-CAAX)* | 5 dpf 2 dpi (7 dpf) | SCI – no bridging | 17 |
| B | *Tg(gfap:EGFP; elavl3:mCherry-CAAX)* | 5 dpf 2 dpi (7 dpf) | SCI – axonal bridging only | 2 |
| B | *Tg(gfap:EGFP; elavl3:mCherry-CAAX)* | 5 dpf 2 dpi (7 dpf) | SCI – glial & axonal bridging | 1 |

| | | | | |
|---|---|---|---|---|
| B | *Tg(gfap:EGFP; elavl3:mCherry-CAAX)* | 5 dpf 3 dpi (8 dpf) | SCI – no bridging | 7 |
| B | *Tg(gfap:EGFP; elavl3:mCherry-CAAX)* | 5 dpf 3 dpi (8 dpf) | SCI – axonal bridging only | 5 |
| B | *Tg(gfap:EGFP; elavl3:mCherry-CAAX)* | 5 dpf 3 dpi (8 dpf) | SCI – glial & axonal bridging | 8 |
| B | *Tg(gfap:EGFP; elavl3:mCherry-CAAX)* | 5 dpf 4 dpi (9 dpf) | SCI – no bridging | 2 |
| B | *Tg(gfap:EGFP; elavl3:mCherry-CAAX)* | 5 dpf 4 dpi (9 dpf) | SCI – axonal bridging only | 5 |
| B | *Tg(gfap:EGFP; elavl3:mCherry-CAAX)* | 5 dpf 4 dpi (9 dpf) | SCI – glial & axonal bridging | 13 |
| B | *Tg(gfap:EGFP; elavl3:mCherry-CAAX)* | 5 dpf 5 dpi (10 dpf) | SCI – no bridging | 2 |
| B | *Tg(gfap:EGFP; elavl3:mCherry-CAAX)* | 5 dpf 5 dpi (10 dpf) | SCI – axonal bridging only | 5 |
| B | *Tg(gfap:EGFP; elavl3:mCherry-CAAX)* | 5 dpf 5 dpi (10 dpf) | SCI – glial & axonal bridging | 13 |
| B | *Tg(gfap:EGFP; elavl3:mCherry-CAAX)* | 7 dpf 0 dpi (7 dpf) | SCI – no bridging | 17 |
| B | *Tg(gfap:EGFP; elavl3:mCherry-CAAX)* | 7 dpf 1 dpi (8 dpf) | SCI – no bridging | 17 |
| B | *Tg(gfap:EGFP; elavl3:mCherry-CAAX)* | 7 dpf 2 dpi (9 dpf) | SCI – no bridging | 13 |
| B | *Tg(gfap:EGFP; elavl3:mCherry-CAAX)* | 7 dpf 2 dpi (9 dpf) | SCI – axonal bridging only | 3 |
| B | *Tg(gfap:EGFP; elavl3:mCherry-CAAX)* | 7 dpf 2 dpi (9 dpf) | SCI – glial & axonal bridging | 1 |
| B | *Tg(gfap:EGFP; elavl3:mCherry-CAAX)* | 7 dpf 3 dpi (10 dpf) | SCI – no bridging | 5 |
| B | *Tg(gfap:EGFP; elavl3:mCherry-CAAX)* | 7 dpf 3 dpi (10 dpf) | SCI – axonal bridging only | 5 |
| B | *Tg(gfap:EGFP; elavl3:mCherry-CAAX)* | 7 dpf 3 dpi (10 dpf) | SCI – glial & axonal bridging | 7 |
| B | *Tg(gfap:EGFP; elavl3:mCherry-CAAX)* | 7 dpf 4 dpi (11 dpf) | SCI – no bridging | 3 |
| B | *Tg(gfap:EGFP; elavl3:mCherry-CAAX)* | 7 dpf 4 dpi (11 dpf) | SCI – axonal bridging only | 3 |
| B | *Tg(gfap:EGFP; elavl3:mCherry-CAAX)* | 7 dpf 4 dpi (11 dpf) | SCI – glial & axonal bridging | 11 |
| B | *Tg(gfap:EGFP; elavl3:mCherry-CAAX)* | 7 dpf 5 dpi (12 dpf) | SCI – no bridging | 3 |
| B | *Tg(gfap:EGFP; elavl3:mCherry-CAAX)* | 7 dpf 5 dpi (12 dpf) | SCI – axonal bridging only | 2 |
| B | *Tg(gfap:EGFP; elavl3:mCherry-CAAX)* | 7 dpf 5 dpi (12 dpf) | SCI – glial & axonal bridging | 12 |

**Table S3. Experimental statistics for recovery of swim behavior** (related to Figures 4-7).

| Figure | Transgenic Line (heterozygous) | Stage | Treatment – Condition | Total number of animals |
|---|---|---|---|---|
| 4-7 Panel A, right | *Tg(gfap:EGFP; elavl3:mCherry-CAAX)* | 3 dpf 5 dpi (8 dpf) | Control | 28 |
| 4-7 Panel A, right | *Tg(gfap:EGFP; elavl3:mCherry-CAAX)* | 3 dpf 5 dpi (8 dpf) | SCI – glial & axonal bridging | 8 |
| 4-7 Panel A, right | *Tg(gfap:EGFP; elavl3:mCherry-CAAX)* | 3 dpf 5 dpi (8 dpf) | SCI – glial bridging only | 2 |
| 4-7 Panel A, right | *Tg(gfap:EGFP; elavl3:mCherry-CAAX)* | 3 dpf 5 dpi (8 dpf) | SCI – axonal bridging only | 7 |
| 4-7 Panel A, right | *Tg(gfap:EGFP; elavl3:mCherry-CAAX)* | 3 dpf 5 dpi (8 dpf) | SCI – no bridging | 11 |
| 4-7 Panel B, right | *Tg(gfap:EGFP; elavl3:mCherry-CAAX)* | 5 dpf 5 dpi (10 dpf) | Control | 29 |
| 4-7 Panel B, right | *Tg(gfap:EGFP; elavl3:mCherry-CAAX)* | 5 dpf 5 dpi (10 dpf) | SCI – glial & axonal bridging | 16 |
| 4-7 Panel B, right | *Tg(gfap:EGFP; elavl3:mCherry-CAAX)* | 5 dpf 5 dpi (10 dpf) | SCI – glial bridging only | 0 |
| 4-7 Panel B, right | *Tg(gfap:EGFP; elavl3:mCherry-CAAX)* | 5 dpf 5 dpi (10 dpf) | SCI – axonal bridging only | 8 |
| 4-7 Panel B, right | *Tg(gfap:EGFP; elavl3:mCherry-CAAX)* | 5 dpf 5 dpi (10 dpf) | SCI – no bridging | 6 |
| 4-7 Panel C, right | *Tg(gfap:EGFP; elavl3:mCherry-CAAX)* | 7 dpf 5 dpi (12 dpf) | Control | 46 |
| 4-7 Panel C, right | *Tg(gfap:EGFP; elavl3:mCherry-CAAX)* | 7 dpf 5 dpi (12 dpf) | SCI – glial & axonal bridging | 10 |
| 4-7 Panel C, right | *Tg(gfap:EGFP; elavl3:mCherry-CAAX)* | 7 dpf 5 dpi (12 dpf) | SCI – glial bridging only | 0 |
| 4-7 Panel C, right | *Tg(gfap:EGFP; elavl3:mCherry-CAAX)* | 7 dpf 5 dpi (12 dpf) | SCI – axonal bridging only | 6 |
| 4-7 Panel C, right | *Tg(gfap:EGFP; elavl3:mCherry-CAAX)* | 7 dpf 5 dpi (12 dpf) | SCI – no bridging | 23 |

**Table S4. Experimental statistics for recovery of swim behavior** (related to Figure 8).

| Panel | Transgenic Line (heterozygous) | Stage | Treatment – Condition | Total number of animals |
|---|---|---|---|---|
| A right | *Tg(gfap:EGFP; elavl3:mCherry-CAAX)* | 3 dpf 7 dpi (10 dpf) | Control | 47 |
| A right | *Tg(gfap:EGFP; elavl3:mCherry-CAAX)* | 3 dpf 7 dpi (10 dpf) | SCI – glial & axonal bridging | 34 |
| A right | *Tg(gfap:EGFP; elavl3:mCherry-CAAX)* | 3 dpf 7 dpi (10 dpf) | SCI – glial bridging only | 0 |
| A right | *Tg(gfap:EGFP; elavl3:mCherry-CAAX)* | 3 dpf 7 dpi (10 dpf) | SCI – axonal bridging only | 18 |
| A right | *Tg(gfap:EGFP; elavl3:mCherry-CAAX)* | 3 dpf 7 dpi (10 dpf) | SCI – no bridging | 8 |
| B right | *Tg(gfap:EGFP; elavl3:mCherry-CAAX)* | 5 dpf 7 dpi (12 dpf) | Control | 29 |
| B right | *Tg(gfap:EGFP; elavl3:mCherry-CAAX)* | 5 dpf 7 dpi (12 dpf) | SCI – glial & axonal bridging | 13 |
| B right | *Tg(gfap:EGFP; elavl3:mCherry-CAAX)* | 5 dpf 7 dpi (12 dpf) | SCI – glial bridging only | 0 |
| B right | *Tg(gfap:EGFP; elavl3:mCherry-CAAX)* | 5 dpf 7 dpi (12 dpf) | SCI – axonal bridging only | 5 |
| B right | *Tg(gfap:EGFP; elavl3:mCherry-CAAX)* | 5 dpf 7 dpi (12 dpf) | SCI – no bridging | 4 |
| C right | *Tg(gfap:EGFP; elavl3:mCherry-CAAX)* | 7 dpf 7 dpi (14 dpf) | Control | 29 |
| C right | *Tg(gfap:EGFP; elavl3:mCherry-CAAX)* | 7 dpf 7 dpi (14 dpf) | SCI – glial & axonal bridging | 10 |
| C right | *Tg(gfap:EGFP; elavl3:mCherry-CAAX)* | 7 dpf 7 dpi (14 dpf) | SCI – glial bridging only | 0 |
| C right | *Tg(gfap:EGFP; elavl3:mCherry-CAAX)* | 7 dpf 7 dpi (14 dpf) | SCI – axonal bridging only | 4 |
| C right | *Tg(gfap:EGFP; elavl3:mCherry-CAAX)* | 7 dpf 7 dpi (14 dpf) | SCI – no bridging | 6 |

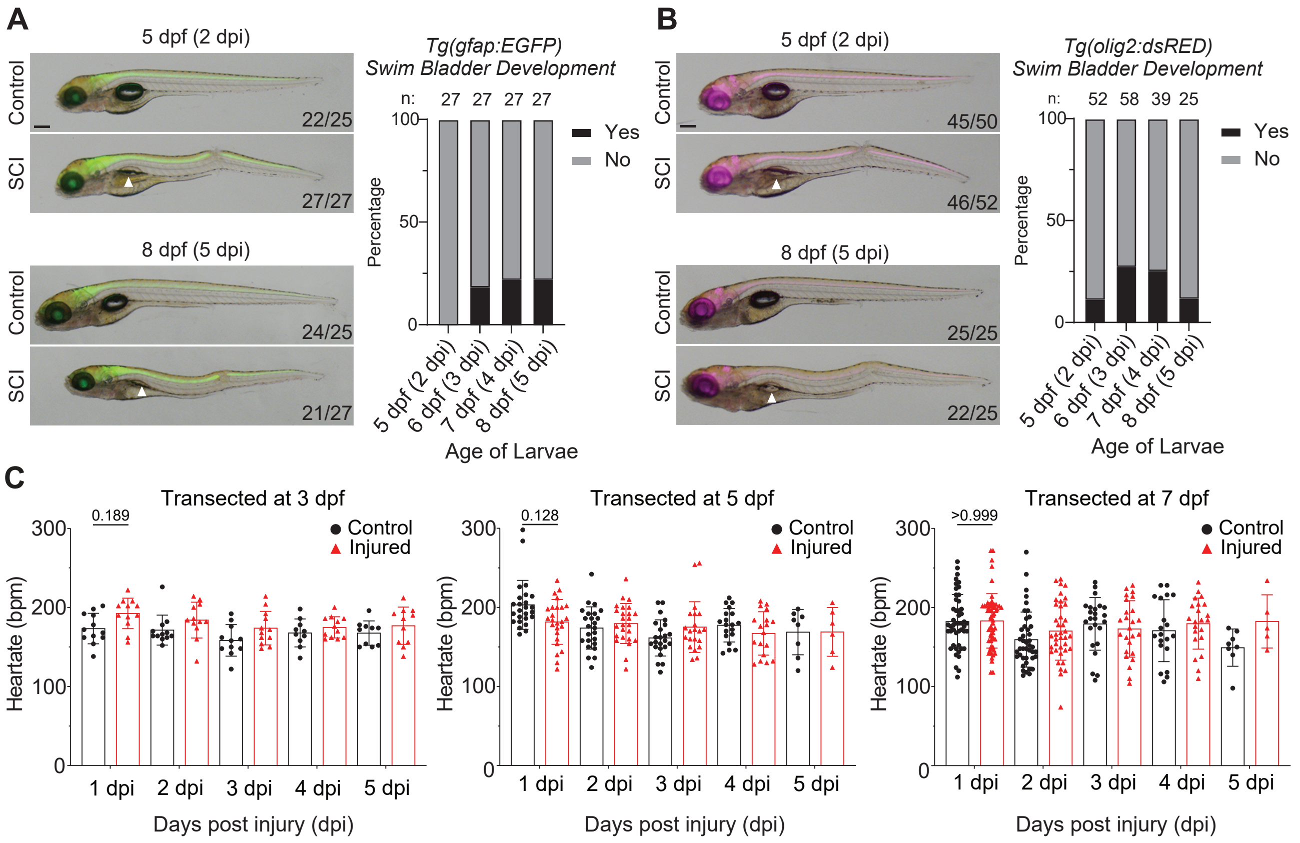

**Figure S1: SCI affects proper swim bladder inflation and heart rate after injury.** (**A**) *Tg(gfap:EGFP)* larvae were transected at 3 dpf and monitored for swim bladder inflation through 5 days post injury (dpi). (**B**) *Tg(Olig2:dsRed)* larvae were transected at 3 dpf and monitored for swim bladder inflation through 5 dpi. The number of larvae with inflated and uninflated (white arrowhead) swim bladders was quantified. Representative micrographs are shown. Larval orientation: lateral view, anterior left. Scale bar = 200 µm. (**C**) Heart rate was quantified by an independent observer each day post injury for transected and control larvae. Individual larvae were included until day of death or until gill development precluded measurement. Control: n=11-23, SCI: initial n=12-25. Individual fish plotted, mean ± SD is shown.

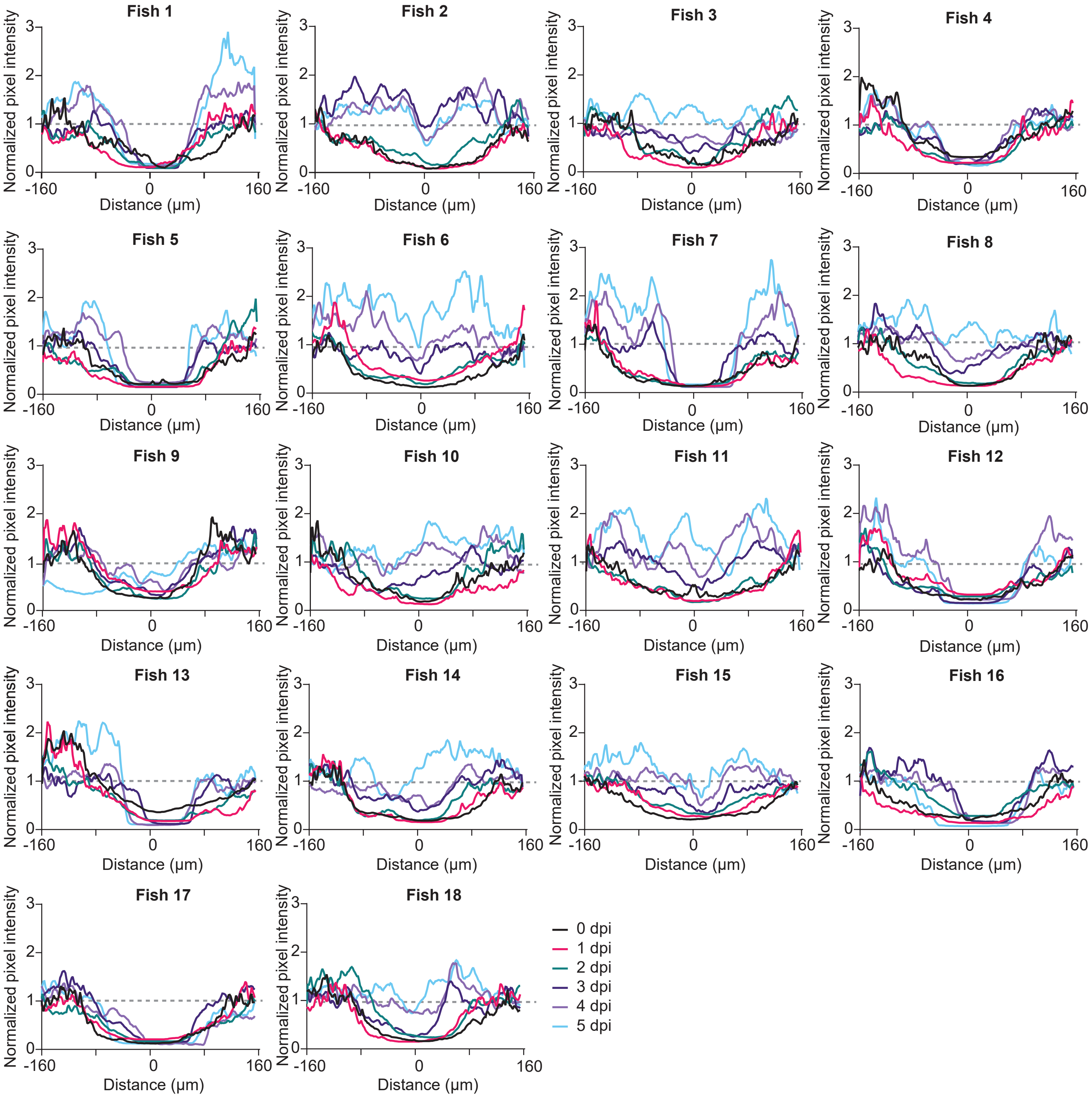

**Figure S2: Glial bridging after SCI in 3 dpf larvae.** (**A**) *Tg(gfap:EGFP; elavl3:mCherry_CAAX)* larvae were transected at 3 dpf and imaged daily. Quantification of normalized mean pixel intensity (solid line) of the GFP signal for each larvae graphed approximately every 10 μm for clarity. Epicenter of the lesion denoted as 0 on the X-axis was determined by the lowest average normalized pixel intensity value on day of injury (0 dpi).

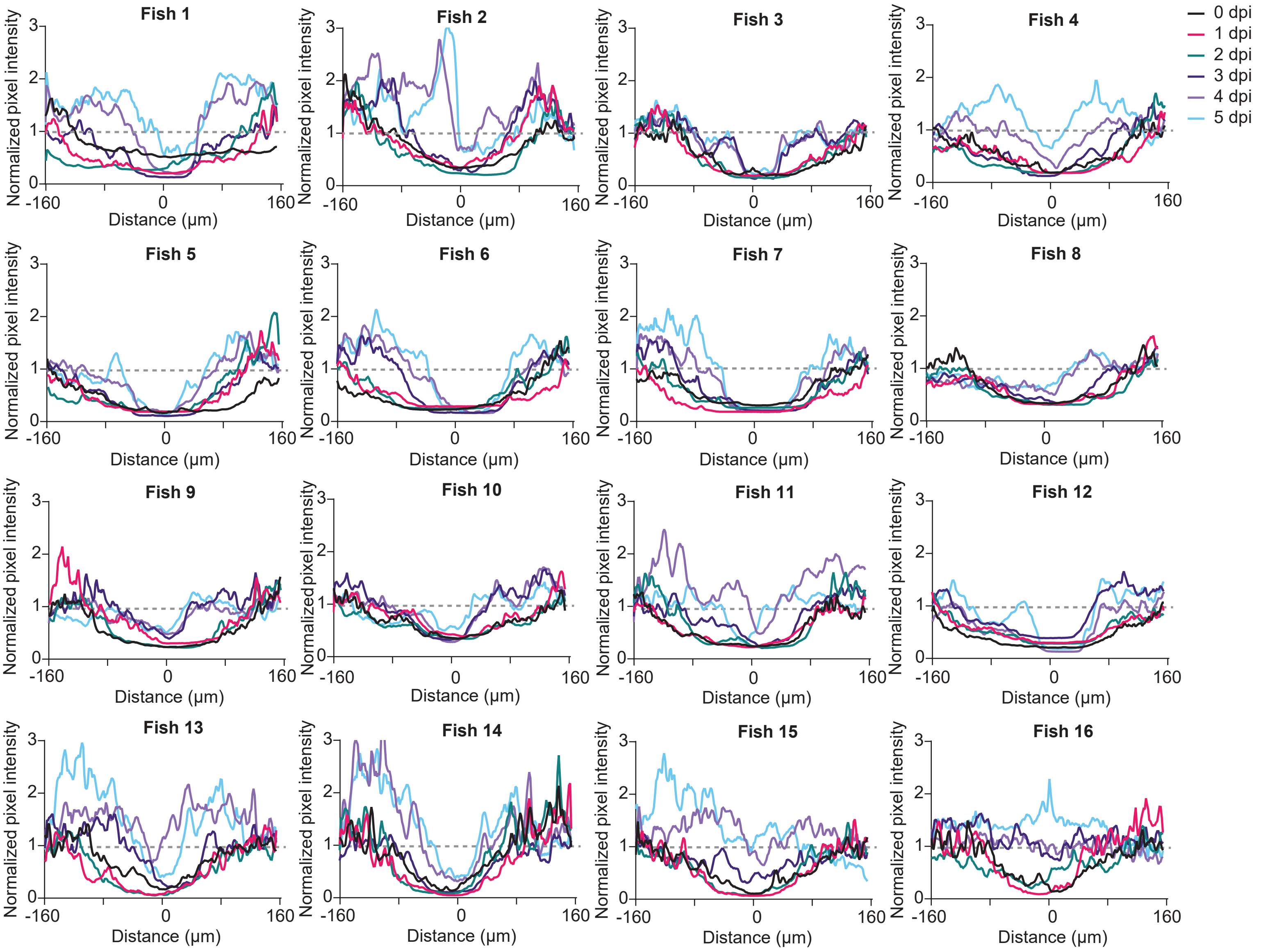

**Figure S3: Glial bridging after SCI in 5 dpf larvae.** (**A**) *Tg(gfap:EGFP; elavl3:mCherry_CAAX)* larvae were transected at 5 dpf and imaged daily. Quantification of normalized mean pixel intensity (solid line) of the GFP signal for each larvae graphed approximately every 10 µm for clarity. Epicenter of the lesion denoted as 0 on the X-axis was determined by the lowest average normalized pixel intensity value on day of injury (0 dpi). GFP signal is higher around the site of injury than in the injury site intself.

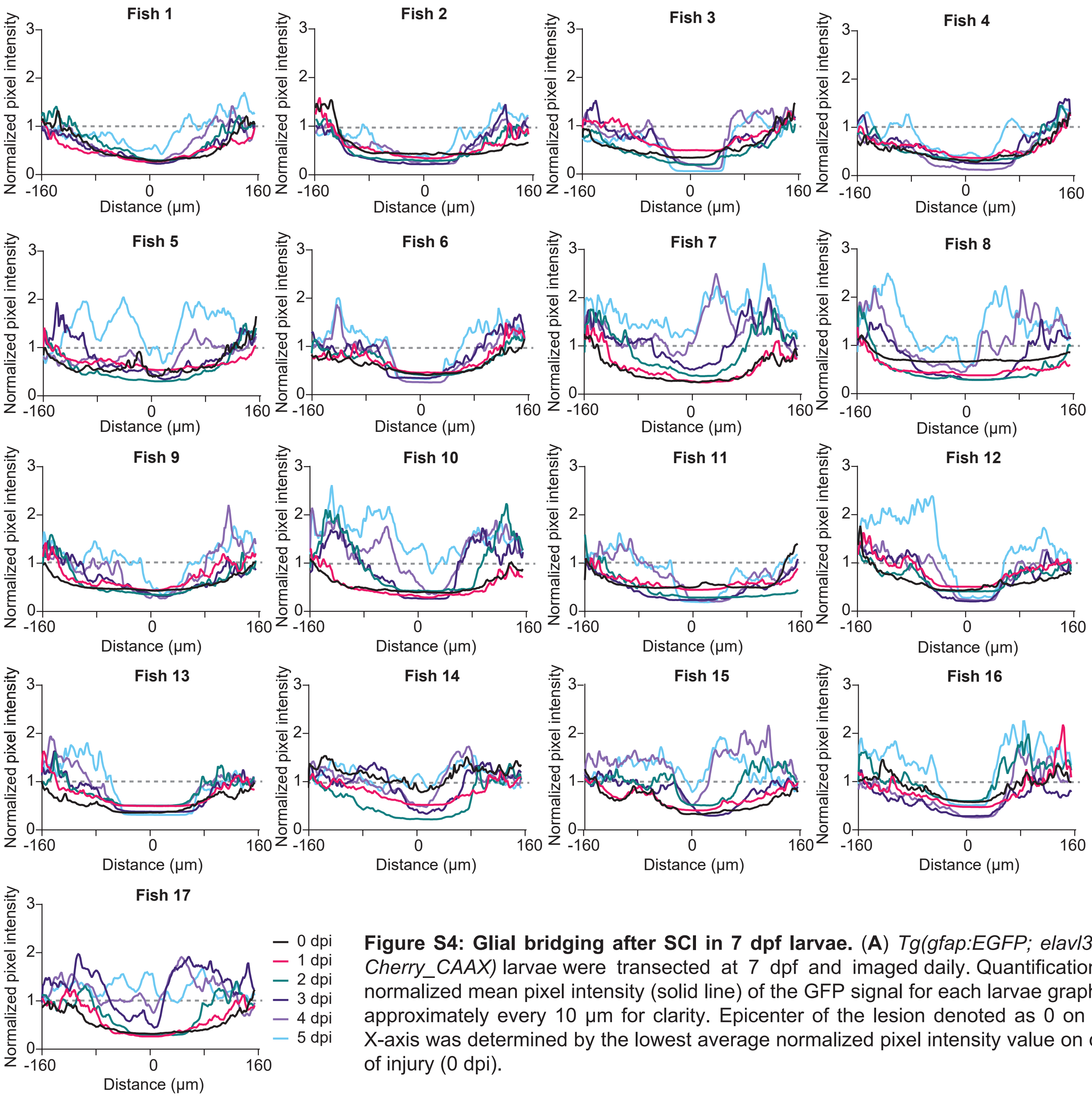

**Figure S4: Glial bridging after SCI in 7 dpf larvae.** (**A**) *Tg(gfap:EGFP; elavl3:m-Cherry_CAAX)* larvae were transected at 7 dpf and imaged daily. Quantification of normalized mean pixel intensity (solid line) of the GFP signal for each larvae graphed approximately every 10 µm for clarity. Epicenter of the lesion denoted as 0 on the X-axis was determined by the lowest average normalized pixel intensity value on day of injury (0 dpi).
